# The *Fusarium graminearum* effector protease FgTPP1 suppresses immune responses and facilitates Fusarium Head Blight Disease

**DOI:** 10.1101/2024.08.30.610543

**Authors:** Martin Darino, Namrata Jaiswal, Reynaldi Darma, Erika Kroll, Martin Urban, Youhuang Xiang, Moumita Srivastava, Hye-Seon Kim, Ariana Myers, Steven R. Scofield, Roger W. Innes, Kim E. Hammond-Kosack, Matthew Helm

## Abstract

Most plant pathogens secrete effector proteins to circumvent host immune responses, thereby promoting pathogen virulence. One such pathogen is the fungus *Fusarium graminearum*, which causes Fusarium Head Blight (FHB) disease on wheat and barley. Transcriptomic analyses revealed that *F. graminearum* expresses many candidate effector proteins during early phases of the infection process, some of which are annotated as proteases. However, the contributions of these proteases to virulence remains poorly defined. Here, we characterize a *F. graminearum* endopeptidase, FgTPP1 (FGSG_11164), that is highly upregulated during wheat spikelet infection and is secreted from fungal cells. To elucidate the potential role of FgTPP1 in *F. graminearum* virulence, we generated *FgTPP1* deletion mutants (*ΔFgtpp1*) and performed FHB infection assays. While the number of completely bleached spikes infected by *F*. *graminearum* wild-type reached 50% of total infected spikes, the number of fully bleached spikes infected by *ΔFgtpp1* mutants was 25%, suggesting FgTPP1 contributes to fungal virulence. Transient expression of green fluorescent protein (GFP)-tagged FgTPP1 revealed that FgTPP1 localizes, in part, to chloroplasts and attenuates chitin-mediated activation of mitogen-activated protein kinase (MAPK) signaling, reactive oxygen species production, and cell death induced by an autoactive disease resistance protein when expressed *in planta*. Notably, the FgTPP1 protein is conserved across the *Ascomycota* phylum, making it a core effector among ascomycete plant pathogens. These properties make FgTPP1 an ideal candidate for decoy substrate engineering, with the goal of engineering resistance to FHB, and likely other crop diseases caused by ascomycete fungi.

## INTRODUCTION

Most fungal phytopathogens express and secrete a repertoire of proteins known as effectors during pathogenesis that modulate plant cell defense responses to facilitate infection and thus disease progression (Bentham et al. 2020). Once secreted, effectors can either be retained in the plant apoplast or translocated directly into host cells where they target multiple host proteins, thereby interfering with host cell-derived defense responses (Selin et al., 2016). To circumvent the immune-modulating activities of effectors, plants have evolved a two-tiered immune signaling network composed primarily of cell surface-localized and intracellular immune receptors (Chen et al. 2022; McCombe et al. 2022; Rhodes et al. 2022). Activation of cell surface-localized immune receptors by pathogen-associated molecular patterns (PAMPs) initiates an intracellular immune signaling cascade that includes, in part, the increased production of extracellular reactive oxygen species (ROS), up-regulation of defense-related gene expression, and activation of mitogen-activated protein kinase (MAPK) signaling (Chen et al. 2022; McCombe et al. 2022; Rhodes et al. 2022). In turn, fungal phytopathogens have evolved effectors that are capable of suppressing ROS generation and accumulation as well as MAPK signaling (Bentham et al. 2020; Chen et al. 2022; Deng et al. 2022; Ngou et al. 2022; Rogers et al. 2024).

Translocated effectors secreted by phytopathogens are diverse in terms of their sizes and specific functions. In the case of bacterial phytopathogens, many effectors have been shown to function as proteases that target specific host proteins to inactivate them (Chandrasekaran et al. 2016; Jashni et al. 2015). For example, the AvrRpt2 protease from *Pseudomonas syringae* is secreted into Arabidopsis host cells, where it cleaves Arabidopsis RPM1-interacting protein 4 (RIN4) protein at two positions (Kim et al. 2005). RIN4 also interacts with proteins in the exocyst complex and appears to regulate callose deposition in response to AvrRpm1 (Kim et al. 2005; Redditt et al. 2019). Similarly, the cysteine protease AvrPphB from *P. syringae* pv. *phaseolicola* targets a family of serine/threonine receptor-like cytoplasmic kinases involved in regulating cell surface-mediated immunity, rendering these kinases inactive (Shao et al. 2003; Zhang et al. 2010). In Arabidopsis, cleavage of one of these kinases, PBS1, by AvrPphB activates the intracellular immune receptor, RPS5, which initiates a signal-transduction cascade culminating in resistance to *P. syringae* strains expressing AvrPphB (Ade et al. 2007; Shao et al. 2003; Simonich and Innes 1995). Importantly, investigating how the Arabidopsis PBS1-RPS5 immune signaling module is activated by the AvrPphB protease has enabled researchers to bioengineer new-to-nature disease resistance specificities against plant pathogens in Arabidopsis as well as crop plants (Carter et al. 2019; Helm et al. 2019; Kim et al. 2016; Pottinger et al. 2020).

The Ascomycete fungal pathogen, *Fusarium graminearum*, causes Fusarium Head Blight (FHB) disease in wheat (*Triticum species*), barley (*Hordeum vulgare*) and other cereal crops, often causing premature senescence and blighting of wheat floral tissues (Dean et al. 2012; Figueroa et al. 2018; Kanja et al. 2021). FHB is considered one of the most economically important fungal diseases of cereal grains, with estimated economic losses exceeding ∼1 billion (USD) annually in direct and indirect effects (Figueroa et al. 2018; Nganje et al. 2004). In addition to reducing overall grain yields, FHB disease contaminates the remaining grain with sesquiterpenoid trichothecene mycotoxins such as deoxynivalenol (DON), which, in turn, affects grain marketability and threatens food safety (Figueroa et al., 2018; Hohn and Desjardins, 1992; Johns et al., 2022). *F. graminearum* employs a hemibiotrophic infection strategy to colonize different host structures such as wheat floral tissues and coleoptiles (Brown et al. 2011; Mentges et al. 2020; Qiu et al. 2019). During the symptomless infection phase, unbranched hyphae spread on the exterior surfaces of the host and form appressoria-like infection cushion structures that enable direct penetration of the cell wall of wheat cells (Mentges et al. 2020; Qiu et al. 2019). Once established, the fungus develops bulbous biotrophic hyphae that branch and push into the host cell displacing the central vacuole (Qiu et al. 2019). Importantly, the plasma membrane of the invaded wheat cells remains intact, indicating that the early phases of infection are biotrophic (Brown et al. 2017; Qiu et al. 2019). During the later combined asymptomatic and symptomatic phases of disease development, *F. graminearum* develops extracellular hyphae to advance within and colonize the apoplastic environment (Brown et al. 2011; Qiu et al. 2019). Coincident with the appearance of disease symptoms, hyphae penetrate into adjacent wheat cells and this change is often accompanied by host cell death (Brown et al. 2011; Qiu et al. 2019). The DON mycotoxin is required for successful traversing of the plasmodesmatal connections between wheat cells (Armer et al. 2024a). Collectively, fungal internal colonization and systemic spread through the wheat rachis results in photobleached spikelets (Brown et al. 2012, 2017).

Similar to other plant pathogenic fungi, *F. graminearum* is predicted to secrete effector proteins during early phases of infection (Brown et al. 2012, 2017; Hao et al. 2020, 2023; Lu et al. 2016; Miltenburg et al. 2022). However, our general knowledge of how *F. graminearum* effector proteins contribute to plant pathogenesis remains limited. Nevertheless, several *F. graminearum* effectors have been identified and reported to have a functional role in fungal pathogenicity. For example, Jiang and colleagues (2020) showed that *F. graminearum* secretes an effector protein designated FgOSP24 (<u>O</u>rphan <u>S</u>ecreted <u>P</u>rotein 24) into the host cell cytoplasm where it subsequently interacts with and promotes the degradation of the SNF1-related kinase, TaSnRK1a. Recent work by Hao et al. (2023) identified a *F. graminearum* effector, FGSG_04563 (FgNls1), that is highly expressed during early phases of FHB disease development, localizes to plant cell nuclei, and interacts with wheat histone 2B (TaH2B). Importantly, deletion of FgNls1 from *F. graminearum* reduced disease progression within wheat spikes and transgenic wheat that silence expression of FgNls1 suppressed FHB disease development, indicating a functional role for FgNls1 in fungal pathogenesis (Hao et al. 2023). Similarly, two candidate *F. graminearum* effectors, ARB93B and FGSG_01831, have been shown to be expressed during the early stages of infection and to suppress cell surface-triggered immune responses, including PAMP-triggered reactive oxygen species production (Hao et al. 2019, 2020).

In addition to the aforementioned *F. graminearum* effectors, two candidate effector proteases have been identified that appear to contribute to *F. graminearum* virulence. Specifically, deletion of a fungalysin metallopeptidase known as FgFly1 (FGSG_03467) and a subtilisin-like protease termed FgPrb1 (FGSG_00192) attenuated *F. graminearum* virulence and FHB disease progression in wheat spikes (Wang et al. 2022; Xu et al. 2020). Recent work by Xiong and colleagues (2024) identified five subtilisin-like proteases from *F. graminearum* (FgSLP1-5) that triggered cell death in *Nicotiana benthamiana*, Arabidopsis, and cotton (*Gossypium barbadense*), and such cell death-inducing activities were independent of BAK1, SOBIR1, EDS1, and PAD4-mediated signaling. Importantly, *F. graminearum* mutants lacking two subtilisin-like proteases, FgSLP1 and FgSLP2, showed reduced fungal virulence in wheat, demonstrating FgSLP1 and FgSLP2 likely contribute to virulence (Xiong et al. 2024). Recently, K. Liu et al. (2024) identified a serine carboxpeptidase, FgSCP (FGSG_08454) that is expressed during early FHB disease development. Deletion of FgSCP in inhibited fungal reproduction, altered vegetative growth, as well as reduced *F. graminearum* virulence and DON biosynthesis in wheat (K. Liu et al. 2024). Collectively, these studies highlight the important roles of *F. graminearum* proteases during pathogenicity. However, despite the importance of *F. graminearum* globally, host disease resistance genes whose protein products recognize the activities of one or more effector proteases from this fungus have yet to be identified. There is thus substantial interest in developing novel methods for conferring resistance to *F. graminearum* via leveraging our understanding of *F. graminearum* virulence mechanisms.

To further understand how candidate secreted effector proteases from *F. graminearum* contribute to fungal virulence we initially took a computational biology approach to identify the predicted secreted proteases present in the most recent *F. graminearum* genome annotation and identified one candidate protease, FGSG_11164 (FGRAMPH1_01G21371), that was highly expressed during floral spike colonization (Brown et al. 2012, 2017; King et al. 2015, 2017a). FGSG_11164 encodes a putative trypsin precursor protein and, therefore, has been designated FgTPP1 (*Fusarium graminearum* <u>T</u>rypsin <u>p</u>recursor <u>p</u>rotein 1). Here, we show that the predicted signal peptide from FgTPP1 is functional and facilitates secretion of FgTPP1 from fungal cells. Importantly, *F. graminearum* mutants lacking FgTPP1 expression (Δ*Fgtpp1*) showed a significantly slower disease progression and colonization of wheat spikes when compared to the wild-type *F. graminearum* PH-1, indicating that FgTPP1 contributes to fungal virulence. Agrobacterium-mediated transient expression of green fluorescent protein (GFP)-tagged FgTPP1 revealed that FgTPP1 localizes, in part, to chloroplast stroma and attenuates chitin-mediated activation of mitogen-activated protein kinase (MAPK) signaling, reactive oxygen species production, and cell death induced by an autoactive disease resistance protein when expressed *in planta*, demonstrating FgTPP1 suppresses multiple plant immune responses. Lastly, we show the TPP1 protein is conserved in fungal species from different genera of the *Ascomycota* phylum suggesting a conserved role(s) for TPP1 in different plant infecting fungi.

## RESULTS

### *F. graminearum* strain PH-1 expresses multiple candidate effector proteases during early wheat spikelet infection

To identify putative effector proteases secreted by *F. graminearum,* we searched the most recent genome annotation of *F. graminearum* (YL1 version, NCBI GenBank number: PRJNA782099) for the terms ‘peptidase’ and ‘protease’ and screened this gene subset for the presence of signal peptides, which would suggest the putative proteases are secreted. This search identified 95 genes encoding putative secreted proteases. To further narrow this list, we selected genes that were upregulated during the early symptomless stages following *F. graminearum* infection of wheat spikes (at 3 and/or 7 days post-inoculation) (Brown et al. 2017). These stringent selection criteria identified seven candidate fungal proteases, of which six had been previously tested and shown not to be essential for virulence, at least when individually mutated (Table 1) (Wang et al. 2022; Xiong et al. 2024; Xu et al. 2020). Hence, we focused on functionally characterizing the remaining candidate protease, FGSG_11164 (FGRAMPH1_01G21371).

**Table 1.** Summary of the candidate effector proteases expressed during early wheat spikelet infection.

| Gene ID | Amino Acids | Signal peptide | Predicted annotation | Reference |
| --- | --- | --- | --- | --- |
| FGSG_00192<br>(FgPrb1) | 533 | Yes (1-15) | Cerevisin precursor protein | Xu et al. 2020 |
| FGSG_00806<br>(FgSLP1) | 413 | Yes (1-20) | Subtilisin-like peptidase | Xu et al. 2020<br>Xiong et al. 2024 |
| FGSG_03315<br>(FgSLP2) | 411 | Yes (1-20) | Subtilisin-like peptidase | Xu et al. 2020<br>Xiong et al. 2024 |
| FGSG_08012 | 390 | Yes (1-18) | Proteinase T precursor protein | Xu et al. 2020 |
| FGSG_11164 | 252 | Yes (1-17) | Trypsin precursor protein | This study |
| FGSG_11472 | 875 | Yes (1-17) | Subtilisin-like peptidase | Xu et al. 2020 |
| FGSG_03467<br>(FgFly1) | 631 | Yes (1-23) | Fungalysin metallopeptidase | Wang et al. 2022 |

Analysis of the FGSG_11164 amino acid sequence with InterProScan indicated that this gene encodes a trypsin precursor protein (Figure 1A). We thus designated FGSG_11164 as *Fusarium graminearum* <u>T</u>rypsin <u>p</u>recursor <u>p</u>rotein 1 (FgTPP1). FgTPP1 encodes a predicted 17-amino acid signal peptide sequence and a trypsin-like serine protease domain (amino acids 27-248) (Figure 1A). Further examination and *in silico* modeling of the FgTPP1 protein using AlphaFold2 showed that the protein has three predicted catalytic active sites, histidine-67 (His67), aspartic acid-112 (Asp112), and serine-208 (Ser208) as well as a putative substrate binding site consisting of aspartic acid-202 (Asp202), serine-224 (Ser224), and glycine-226 (Gly226) (Figure 1A and 1B). Intriguingly, FgTPP1 is also predicted to contain a chloroplast targeting peptide sequence (amino acids 62-102), suggesting it may localize to chloroplasts (Figure 1A and 1B).

**Figure 1.**
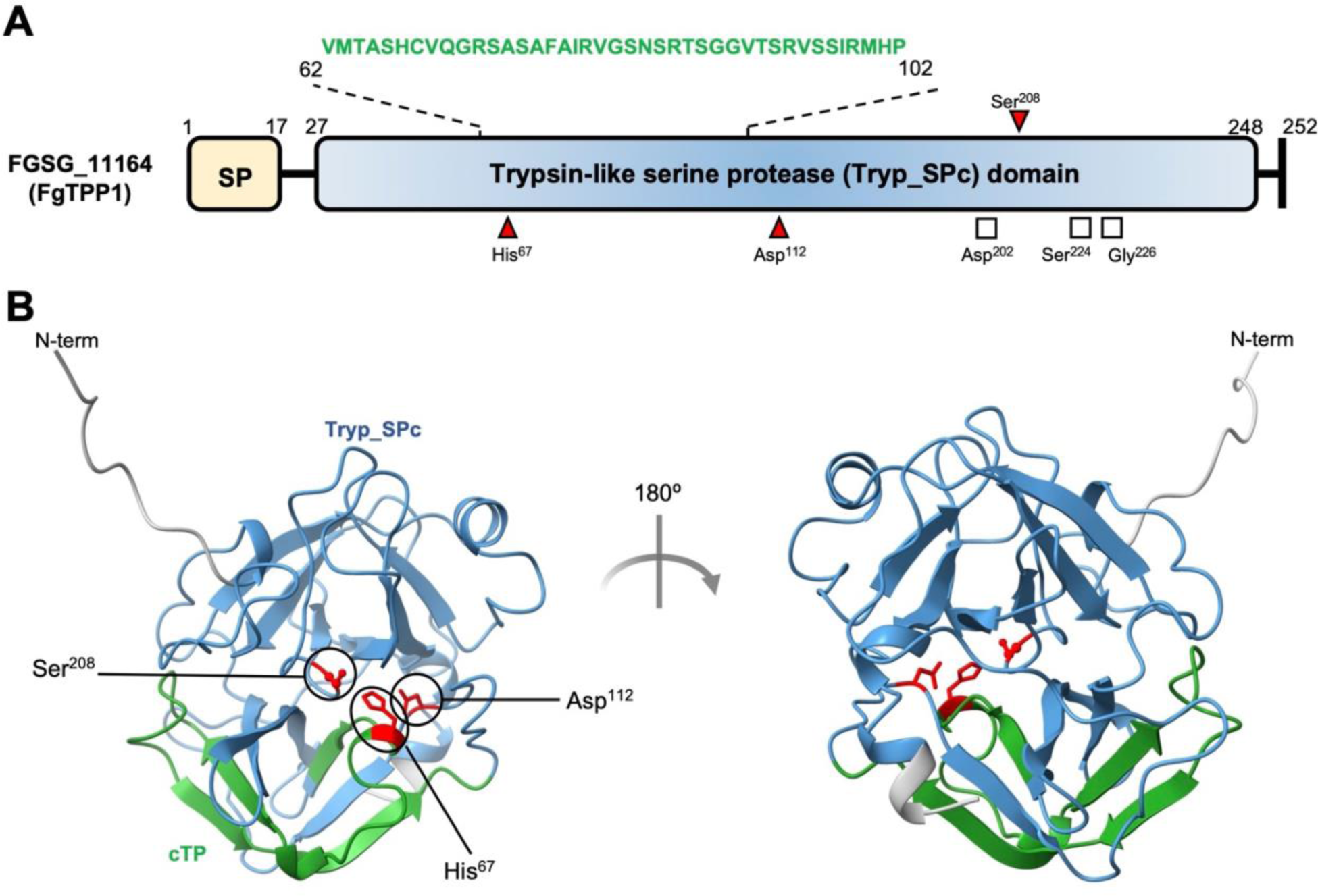
Schematic illustration and predicted protein structure of FgTPP1 (FGSG_11164). **A,** Schematic representation of the *F. graminearum* candidate effector protease FgTPP1 including the putative signal peptide (SP; aa 1-17) sequence and the predicted trypsin-like serine protease (Tryp_SPc; aa 27-248) domain. The predicted chloroplast targeting motif (aa 62-102) is indicated in green. Putative catalytic active sites (His-67, Asp-112, and Ser-208) are indicated by red triangles and predicted substrate binding sites (Asp-202, Ser-224, and Gly-226) are indicated by white squares. Numbers delineate amino acid positions. **B,** The predicted three-dimensional protein structure of FgTPP1 as determined by AlphaFold2. The predicted trypsin-like serine protease domain is indicated in light blue, the chloroplast targeting motif is indicated in green, and the putative catalytic residues are indicated in red. N-term: amino terminus.

### The predicted signal peptide of FgTPP1 confers secretion in yeast cells

To test whether the predicted signal peptide of FgTPP1 is functional, we performed a yeast secretion trap assay (Zhou et al. 2020). This assay employs a *suc2* yeast mutant that is unable to grow on medium with sucrose as the sole carbon source (Zhou et al. 2020). We fused full length *FgTPP1*, including its signal peptide sequence, to a truncated *SUC2* gene lacking a signal peptide (SUC2^22-511^), generating a pGAD-FgTPP1:SUC2^22–511^ construct, which was subsequently transformed into a *suc2* yeast mutant. As a negative control, the truncated *SUC2* gene was fused to *FgTPP1* lacking its signal peptide sequence (pGAD-FgTPP1^ΔSP^:SUC2^22–511^). A pGAD-SUC2^22–511^ only construct was used as an additional negative control. As a positive control, we fused the *F. graminearum* effector FgOSP24, which was previously shown to be secreted (Jiang et al. 2020), to the truncated SUC2^22–511^ gene and transformed this pGAD-FgOSP24:SUC2^22–511^ construct into a *suc2* yeast mutant. As expected, *suc2* yeast mutants expressing either pGAD-FgTPP1^ΔSP^:SUC2^22–511^ or pGAD-SUC2^22–511^ were unable to grow on yeast synthetic dropout media (SD) supplemented with sucrose as the sole carbon source, indicating FgTPP1^ΔSP^ and SUC2 proteins are not secreted (Figure 2). Consistent with our hypothesis, *suc2* yeast mutants transformed with either pGAD-FgTPP1:SUC2^22–511^ or pGAD-FgOSP24:SUC2^22–511^ consistently grew on SD media supplemented with glucose or sucrose, demonstrating that the FgOSP24-SUC2 and FgTPP1-SUC2 fusion proteins are secreted (Figure 2). These data thus confirm that the predicted signal peptide from FgTPP1 is indeed functional and confers secretion of FgTPP1 from fungal cells.

**Figure 2.**
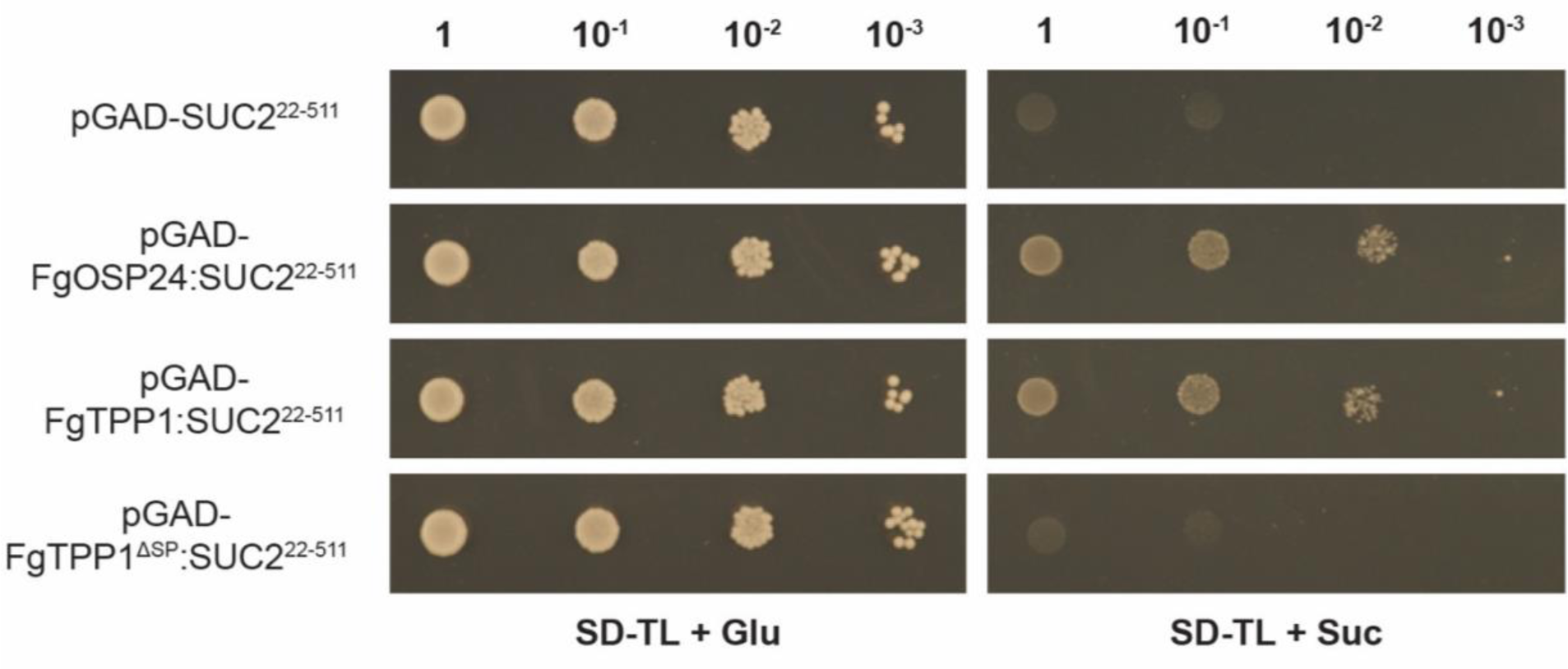
The FgTPP1 protein encodes a functional secretion signal. A yeast secretion trap assay was performed to test functionality of the predicted signal peptide from FgTPP1 (Zhou et al. 2020). A yeast strain lacking the invertase SUC2 was transformed with pGAD-*FgTPP1*:SUC2^22–511^, which contains a truncated SUC2 gene, without its signal peptide (SUC2^22-511^) fused with full length *FgTPP1*, including its signal peptide sequence, or pGAD-*FgTPP1^ΔSP^*:SUC2^22–511^, which lacks the predicted signal peptide sequence from *FgTPP1*. As a positive control, the *suc2* yeast mutant was transformed with pGAD-*FgOSP24*:SUC2^22–511^, which contains the full length *FgOSP24* effector from *F. graminearum* and which was previously shown to be secreted from fungal cells (Jiang et al. 2020). The pGAD-*FgTPP1^ΔSP^*:SUC2^22–511^ and pGAD-SUC2^22–511^ constructs were used as negative controls. Synthetic dropout media (SD) lacking tryptophan and leucine (-TL) supplemented with 2% glucose (Glu) was used as a control media. Images were taken after 4 days of growth and two independent replicates were performed with similar results.

### FgTPP1 contributes to fungal virulence

Transcriptome data sets obtained from detailed spatial and temporal analyses of the early symptomatic and asymptomatic phases of *F. graminearum* colonization of wheat spikes revealed that the *FgTPP1* gene is strongly induced during the first week of infection (Brown et al. 2017). To assess whether FgTPP1 contributes to fungal virulence, we used a homologous recombination strategy to replace the *FgTPP1* gene with a cassette expressing a bacterial gene conferring resistance to hygromycin B (Supplementary Figure S1A) (Catlett et al. 2003; King et al. 2017b). Primer combination P20 / P21 confirmed deletion of the *FgTPP1* coding sequence in two independent mutant strains (PH-1-Δ*Fgtpp1-1* and PH-1-Δ*Fgtpp1-3*) whilst primer combinations P16 / P17 and P18 / P19 confirmed replacement of the *FgTPP1* coding sequence by the hygromycin expression cassette (Supplementary Figure S1A and Supplementary Table S4). Both Δ*Fgtpp1* mutants displayed no observable defects in fungal morphology or radial growth when grown on Potato Dextrose Agar (PDA) plates, including under different stress conditions (Supplementary Figure S1B). Finally, no noticeable defects were observed in perithecia formation and DON mycotoxin production between the PH-1-Δ*Fgtpp1-1* mutant and wild-type *F. graminearum* PH-1 strain (Supplementary Figure S1C and S1D).

Next, we investigated whether FgTPP1 is involved in the infection process by performing fungal virulence assays using a top inoculation approach. In this assay, fungal spores of the PH-1-Δ*Fgtpp1-1* mutant and wild-type *F. graminearum* PH-1 strain were inoculated into the 5^th^ and 6^th^ spikelets from the top of wheat spikes at anthesis of the susceptible wheat cultivar Bobwhite. Bleached spikelets below the point of inoculation were evaluated 12 days post-infection (dpi). The PH-1-Δ*Fgtpp1-1* mutant was able to infect the inoculated spikelets and systemically spread through the spike at a rate similar to wild-type *F. graminearum* PH-1 (Figure 3A). Consistent with the FHB symptoms, no statistically significant differences in the area under the disease progress curve (AUDPC) were observed between the PH-1-Δ*Fgtpp1-1* mutant and wild-type *F. graminearum* PH-1 strain (Figure 3A). To explore further potential virulence defects in the PH-1-Δ*Fgtpp1-1* mutant, we performed a second inoculation approach, which we refer to as the bottom inoculation assay. In brief, spores from either the PH-1-Δ*Fgtpp1-1,* PH-1-Δ*Fgtpp1-3*, or wild-type *F. graminearum* PH-1 were inoculated into the first two full-sized spikelets located at the base of a wheat spike at anthesis, and completely as well as partially bleached spikes were recorded at 10 dpi. In this approach, photobleaching of the spikelets may be a consequence of fungal spread and/or obstruction of the plant vascular tissue, which, in turn, may lead to cell death in wheat tissues above the inoculation point (Bai and Shaner 2004).). The number of completely bleached spikes inoculated with the wild-type *F. graminearum* PH-1 strain was similar to the number of partially bleached spikes (Figure 3B). Furthermore, the frequencies of partially and fully bleached spikes fitted a 1:1 ratio (Figure 3B). However, in these bottom inoculations, both the PH-1-Δ*Fgtpp1-1* and PH-1-Δ*Fgtpp1-3* mutant strains showed a reduced number of fully bleached spikes. The frequencies of fully and partially bleached spikes for both mutants did not fit a 1:1 ratio as observed for PH-1 (Figure 3B). The reduced number of fully bleached spikes in both mutants indicated a slower FHB disease progression when compared to wild-type PH-1 strain (Figure 3B and Supplementary Figure 2).

**Figure 3.**
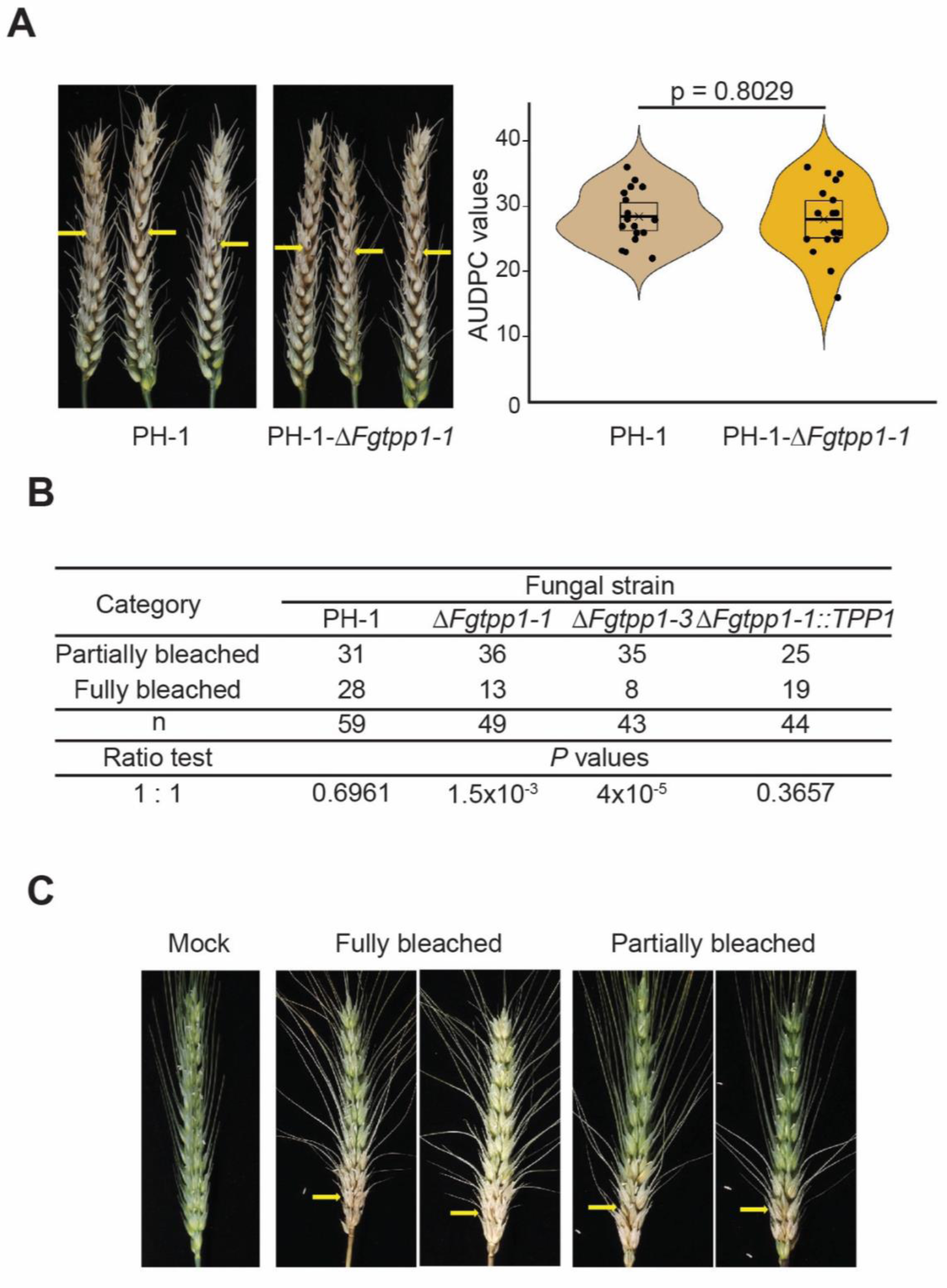
Deletion of the *FgTPP1* gene reduces fungal virulence in bottom inoculated wheat spikes. **A,** The PH-1-Δ*Fgtpp1-1* mutant did not show an observable virulence defect when wheat spikes were inoculated using the top inoculation method. Yellow arrows indicate inoculation points. Wheat spikes infected either with the Δ*Fgtpp1-1* mutant (PH-1-Δ*Fgtpp1-1*) or wild-type *F. graminearum* PH-1 strain showed similar disease symptoms (left) and the AUDPC values calculated for the Δ*Fgtpp1-1* mutant and wild-type PH-1 did not reveal significant differences (right). The statistical analysis (t-test) included pooled data from three independent replicates (n = 43 and 59 infected plants with Δ*Fgtpp1-1* and wild-type *F. graminearum* PH-1 strain, respectively). Violin plots show distribution of the AUDPC values (black dots), average AUDPC values and confidence intervals (rectangles) for each fungal strain. **B,** The Δ*Fgtpp1* mutants (PH-1-Δ*Fgtpp1-1* and PH-1-Δ*Fgtpp1-3*) showed a virulence defect with the bottom inoculation method. The observed number of fully and partially bleached spikes fits to a 1:1 ratio for wild-type *F. graminearum* PH-1 strain whilst the observed frequencies for the Δ*Fgtpp1* mutants deviated significantly from the 1:1 ratio (p ˂ 0.01). Genetic complementation of the mutant strain Δ*Fgtpp1-1* with a full copy of the *Fgtpp1* gene (PH-1-Δ*Fgtpp1-1::TPP1*) restored the virulence defect as the complemented strain fits a 1:1 ratio, which is identical to the wild-type *F. graminearum* PH-1 strain. The statistical analysis included pooled data from four independent replicates. **C,** Representative wheat spike images from the bottom inoculation method at 10 days post-inoculation. Yellow arrows indicate inoculation points. Fully bleached spikes consist of bleached and light green spikelets with curved awns along the entire spike. Partially bleached spikes consist of bleached spikelets with curved awns only in the bottom of the spike whilst the upper spikelets remain dark green with straight awns as observed in the mock control.

To confirm that the observed reduction in FHB disease progression in the bottom inoculation assay was a result of the absence of FgTPP1, the Δ*Fgtpp1-1* mutant was complemented with a copy of full-length *FgTPP1* gene, including its native promoter and terminator, thereby generating the complemented strain PH-1-*ΔFgtpp1-1*::*TPP1*. The complementing copy of *FgTPP1* was inserted a recently described neutral locus in *F. graminearum* namely Target Site Integration (TSI) locus 1 (Darino et al. 2024). The TSI locus 1 allows target site integration of different cassettes without affecting either fungal virulence or fungal growth under different conditions. PCR amplification confirmed the insertion of a single copy of the cassette as primer combination P28 / P29 amplified the expected 6,123 base pair PCR product (Supplementary Figure S1E and Supplementary Table S4). The PH-1-*ΔFgtpp1-1*::*TPP1* strain did not show observable defects in radial growth or fungal morphology when grown under different stress conditions and, importantly, restored the defect in virulence observed with the PH-1-Δ*Fgtpp1-1* mutant (Figure 3B and Supplementary Figures 1B and 2). Collectively, these data demonstrate that FgTPP1 contributes to *F. graminearum* virulence in wheat spikes.

### FgTPP1 accumulates within the stroma of chloroplasts

Analysis of the FgTPP1 amino acid sequence using the deep learning program LOCALIZER (Sperschneider et al. 2017) revealed that FgTPP1 encodes a predicted chloroplast targeting sequence (CTS, amino acids 62-102). To test whether FgTPP1 is indeed targeted to chloroplasts, we fused FgTPP1 (without its signal peptide) to the N-terminus of Green Fluorescent Protein (GFP) and transiently co-expressed it in *Nicotiana benthamiana* with mCherry-tagged Rubisco small subunit transit peptide (RbcS-TP:mCherry), a chloroplast stroma marker protein (Helm et al. 2022; Nelson et al. 2007). Live-cell imaging using laser-scanning confocal microscopy of *N. benthamiana* epidermal cells revealed that FgTPP1:GFP partially localized to the chloroplast stroma as indicated by an overlap in the fluorescence signal from FgTPP1:GFP and RbcS-TP:mCherry (Figure 4A). As a control, we expressed free GFP with RbcS-TP:mCherry to exclude the possibility that the apparent chloroplast localization of FgTPP1:GFP was a result of free GFP or chlorophyll autofluorescence. The experiments showed that the fluorescence signal from free GFP did not significantly overlap with the fluorescence signal from the RbcS-TP:mCherry construct (Figure 4A). We next performed immunoblot analyses to assess protein accumulation and confirm the integrity of the FgTPP1:GFP protein. Intriguingly, these experiments revealed that the FgTPP1:GFP fusion protein consistently produced multiple, distinct protein products, suggesting that the fusion protein may be cleaved prior to or following entry into chloroplasts (Figure 4B). To further confirm chloroplast localization of FgTPP1, we fused the mCherry fluorescent protein to FgTPP1 (FgTPP1:mCherry) and performed live-cell imaging and immunoblot analyses on chloroplasts isolated from *N. benthamiana*. Confocal microscopy imaging of isolated chloroplasts revealed mCherry fluorescence signal in some, but not all, chloroplasts (Figure 4C). Taken together, our results demonstrate that FgTPP1 partially localizes to chloroplasts when transiently expressed in *N. benthamiana*.

**Figure 4.**
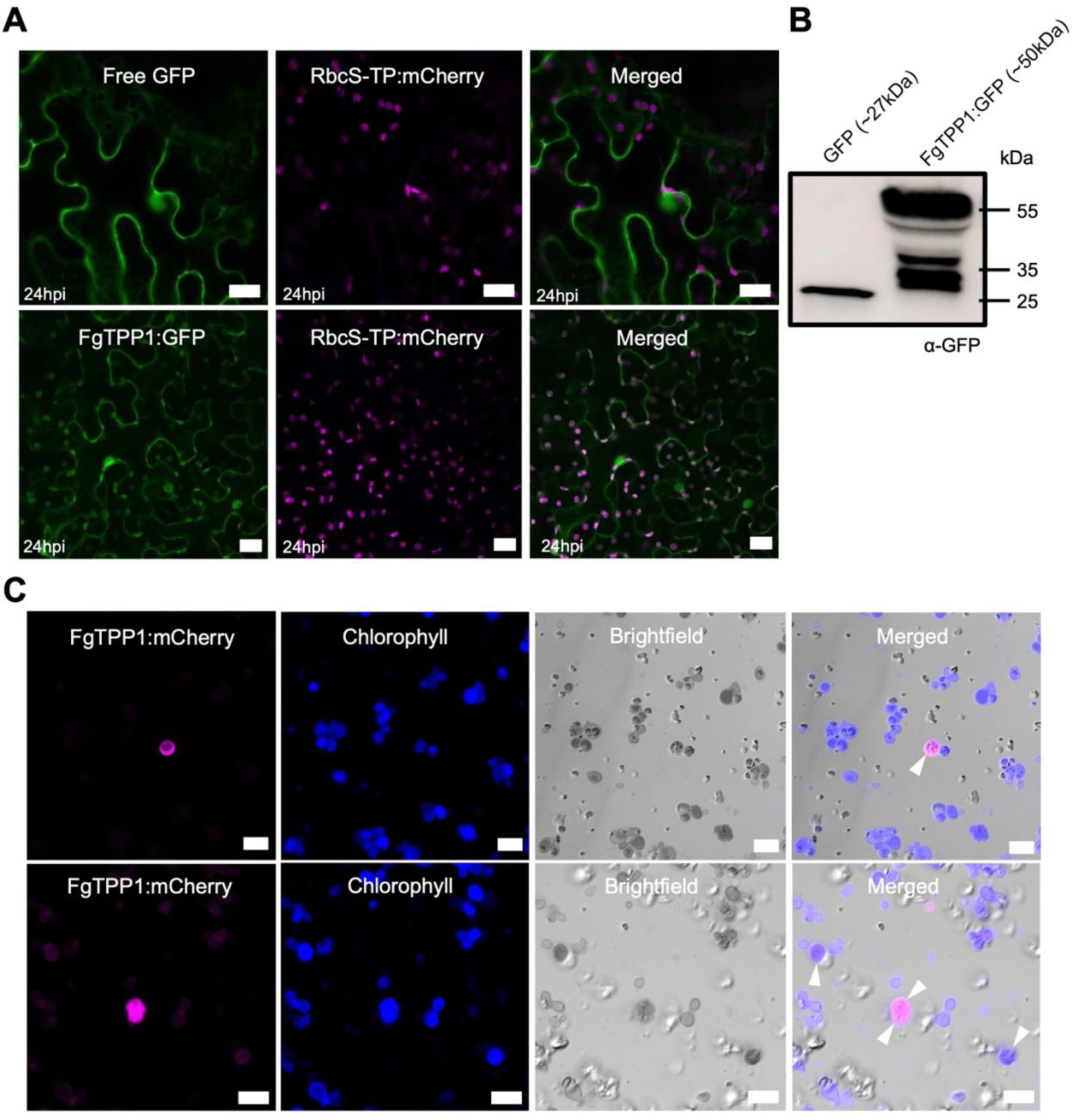
The FgTPP1:GFP fusion protein targets the chloroplast stroma and nucleo-cytosol when transiently expressed in *N. benthamiana*. **A,** Green Fluorescent Protein (GFP)-tagged FgTPP1 (FgTPP1:GFP) localizes to the chloroplast stroma as well as the nucleo-cytosol in *N. benthamiana* epidermal cells. FgTPP1:GFP or free GFP were transiently co-expressed with mCherry-tagged RbcS-TP (RbcS-TP:mCherry) in *N. benthamiana* leaves using *Agrobacterium*-mediated transformation (agroinfiltration). Images were taken 24 hours post-agroinfiltration (hpi). mCherry-tagged RbcS-TP was included as a reference for chloroplast localization (Helm et al. 2022). All confocal micrographs shown are of single optical sections. The scale bar represents 20 microns. **B,** Immunoblot analysis of the FgTPP1:GFP fluorescent protein fusion. The indicated constructs were transiently expressed in *N. benthamiana* leaves. Total protein was isolated 24 hours following agroinfiltration and analyzed using immunoblotting. **C,** mCherry-tagged FgTPP1 localizes to chloroplasts isolated from agroinfiltrated *N. benthamiana*. Leaf tissue transiently expressing FgTPP1:mCherry were harvested 24 hours following agroinfiltration and homogenized in cold isolation buffer. Chloroplasts were isolated as described in the Materials and Methods and imaged using laser-scanning confocal microscopy. Chlorophyll autofluorescence (in blue) was used as a marker for chloroplasts. White arrowheads indicate overlapping mCherry and chlorophyll fluorescence signals. Confocal micrographs from two independent biological replicates are shown in the top and bottom panels. The scale bar shown represents 10 microns.

### FgTPP1 suppresses cell surface- and intracellular-mediated immune responses

Most pathogen effectors function, at least in part, to suppress host immune responses. Prior to testing whether FgTPP1 can suppress immune signaling, we first confirmed that chitin enhances the accumulation of Mitogen-activated protein kinase (MAPK) protein, specifically the phosphorylation of MAPK3 and MAPK6, which has been linked to immune response signaling downstream of pathogen recognition (Jaiswal et al. 2022). *N. benthamiana* leaves transiently expressing free GFP were infiltrated with either deionized water (mock) or chitin, and at 0-, 5-, and 10-minutes following treatment, accumulation of phosphorylated MAPK3 and MAPK6 protein was assessed using immunoblot analyses with MAPK3/6 phospho-specific antibodies These experiments revealed an increase in phosphorylated MAPK3 and MAPK6 protein in response to chitin treatment when compared to the mock control, demonstrating that the increased accumulation of phosphorylated MAPK protein is indeed chitin-mediated (Figure 5A). To test whether FgTPP1 could attenuate chitin-mediated MAPK signaling, we treated *N. benthamiana* leaves transiently expressing either free GFP or FgTPP1:GFP with chitin and assessed accumulation of phosphorylated MAPK3 and MAPK6 protein. These experiments revealed that expression of FgTPP1:GFP protein consistently suppressed the accumulation of phosphorylated MAPK3 and MAPK6 in response to chitin relative to the free GFP control (Figure 5B).

**Figure 5.**
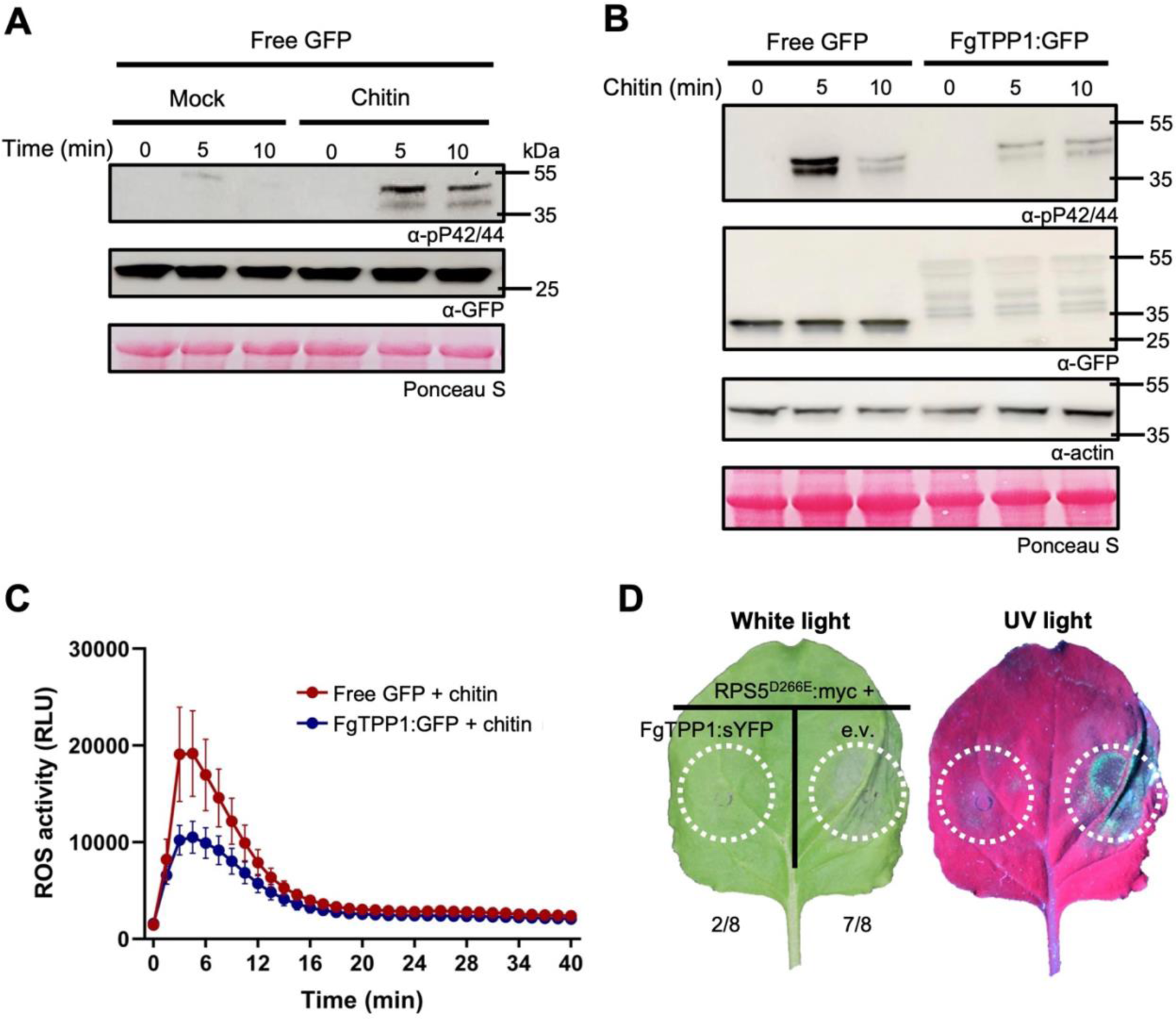
FgTPP1 suppresses cell surface-trigged and intracellular-mediated immune responses. **A,** MAPK3 and MAPK6 phosphorylation is induced upon chitin treatment. *N. benthamiana* leaves transiently expressing free GFP (to control for the presence of *A. tumefaciens*) were agroinfiltrated using a needless syringe. Forty-eight hours following agroinfiltration, leaves were infiltrated with chitin (5 μg/mL chitin [hexamer]) and leaf discs (10 mm diameter) harvested at the indicated time points for protein extraction. Anti-phospho-P42/44 antibodies were used to detect phosphorylated MAPK3 and MAPK6. Mock treated leaves were infiltrated with deionized water. Ponceau S staining was used as a loading control. **B,** Chitin-induced accumulation of phosphorylated MAPK3/6 is attenuated by FgTPP1. The indicated constructs were transiently expressed in *N. benthamiana.* Forty-eight hours after agroinfiltration, leaves were infiltrated with chitin (5 μg/mL chitin [hexamer]), total protein extracted at the indicated time points, and immunoblotted using the indicated antibodies. *N. benthamiana* actin and Ponceau S staining were used as loading controls. **C,** Transient expression of FgTPP1:GFP attenuates chitin-induced ROS burst in *N. benthamiana*. FgTPP1:GFP or free GFP were transiently expressed in *N. benthamiana*. Forty-eight hours after agroinfiltration, *N. benthamiana* leaf discs (5 mm diameter) were collected, treated with chitin (5 μg/mL chitin [hexamer]), and relative luminescence (RLU) was monitored for 40 minutes using a previously optimized luminol-based assay (Rogers et al. 2024). **D,** Suppression of RPS5^D266E^-mediated cell death by FgTPP1 in the *N. benthamiana* leaves. *N. benthamiana* leaves were co-infiltrated with *A. tumefaciens* strains carrying myc-tagged RPS5^D266E^ (OD_600_ = 0.15), FgTPP1:sYFP (OD_600_ = 0.3), or empty vector (e.v.; OD_600_ = 0.3). Forty-eight hours following agroinfiltration, *N. benthamiana* leaves were sprayed with 50 μM dexamethasone to induce protein expression. *N. benthamiana* leaves were assessed for cell death and photographed under white and ultraviolet (UV) light 16 hours post-transgene induction. Fractions indicate the number of leaves with observable HR-like cell death / the total number of leaves agroinfiltrated. Experiments were performed three independent times with similar results.

Several effectors from *F. graminearum* have been previously shown to suppress chitin-triggered reactive oxygen species (ROS) production (Hao et al. 2019, 2020). To test whether FgTPP1 could also suppress chitin-mediated ROS accumulation, we transiently expressed either free GFP or FgTPP1:GFP in *N. benthamiana* leaves. Plants were subsequently challenged with chitin and ROS production was monitored over time using a previously optimized luminol-based assay (Rogers et al. 2024). Expression of FgTPP1:GFP consistently attenuated chitin-mediated ROS accumulation when compared to the free GFP control (Figure 5C). Taken together, our results indicate that FgTPP1 suppresses multiple chitin-mediated immune responses.

Two previously studied effector proteins from *F. graminearum*, namely FgOSP24 and FgNLS1, have been reported to suppress BAX (Bcl-associated X)-induced cell death in *N. benthamiana*, and thus likely have a functional role in subverting host immune responses (Hao et al. 2023; Jiang et al. 2020). To test whether FgTPP1 can suppress cell death, we used an allele of the Arabidopsis disease resistance protein RPS5, RPS5^D266E^, which constitutively activates cell death when transiently expressed in *N. benthamiana* (Ade et al. 2007). We fused FgTPP1 to the N-terminus of super Yellow Fluorescent Protein (sYFP) and cloned FgTPP1:sYFP into the dexamethasone-inducible expression plasmid, pTA7001 (Qi et al. 2012; Vinatzer et al. 2006). We then transiently co-expressed RPS5^D266E^ with either FgTPP1:sYFP or empty vector (e.v.), and assessed for the suppression of RPS5^D266E^-mediated cell death 16 hours post-dexamethasone induction. These experiments revealed that FgTPP1:sYFP consistently attenuated RPS5^D266E^-triggered cell death when compared to the empty vector control (Figure 5D). Collectively, our data demonstrate that FgTPP1 attenuates cell surface-triggered and intracellular-mediated immune responses. Furthermore, the observation that FgTPP1 is secreted, localizes, in part, to chloroplasts, and suppresses immune responses when expressed inside plant cells indicates that FgTPP1 functions inside plant cells and may thus be translocated from the fungus into the host cell during infection.

### *TPP1* alleles and TPP1 protein haplotypes are conserved among plant infecting fungi of the *Ascomycota* phylum

To identify alleles of the *FgTPP1* gene in the *F. graminearum* global population, the nucleotide coding sequence of *FgTPP1* from the *F. graminearum* PH-1 strain was aligned to the nucleotide coding sequences of 27 *F. graminearum* isolates collected in 15 countries covering six continents (Supplementary Table S1). Seven alleles of the *FgTPP1* gene were identified (alleles 1 to 7) (Figure 6A). However, only two protein haplotypes were identified from these 7 alleles (Figure 6). Protein haplotype I is encoded by alleles 1 to 5 whilst protein haplotype F is encoded by alleles 6 and 7. The difference between the two protein haplotypes resides in amino acid residue 4, where haplotype I possesses an isoleucine whilst haplotype F possesses phenylalanine (Figure 6B). This amino acid change resides within the secretion signal, but both protein haplotypes are predicted to be secreted as determined by SignalP v6.0 (Teufel et al. 2022). Approximately 85.7% of *F. graminearum* strains, including PH-1, have the protein haplotype I whereas only four recent European strains collected from western bordering countries, including France, Belgium, and Luxembourg, between 2002 and 2008 have protein haplotype F. Taken together, our analyses revealed the TPP1 protein sequence is highly conserved among *F. graminearum* isolates collected from diverse geographic regions during the last 50 years.

**Figure 6.**
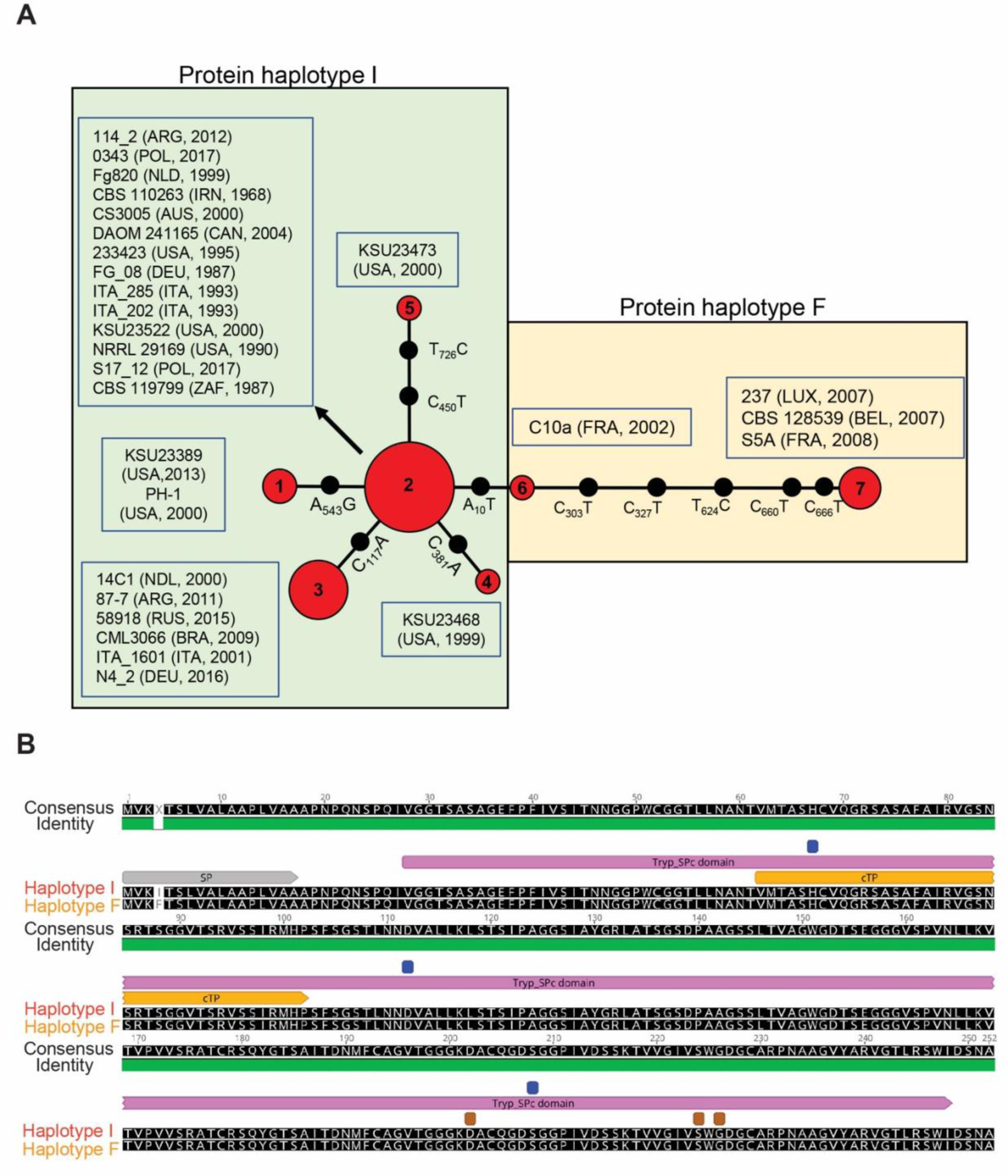
*TPP1* alleles and TPP1 protein haplotypes in *F. graminearum* isolates collected globally. **A,** Allele networks for the *FgTPP1* coding sequences from 28 *F. graminearum* isolates collected worldwide. Red circles represent one allele and the number within each of the red circles represents the allele number. Each black circle represents the number of mutations between adjacent alleles. The mutated bases in alleles 2 to 7 are compared to allele 1. Five alleles (alleles 1 to 5) encode the protein haplotype isoleucine (I) whilst two alleles (alleles 6 and 7) encode protein haplotype phenylalanine (F). Haplotype networks were generated in POPart using a minimum spanning network. The country of origin and year where each isolate was collected are given between brackets. ARG, Argentina; AUS, Australia; BEL, Belgium, BRA, Brazil; CAN, Canada; FRA, France; DEU, Germany; ITA, Italy; IRN, Iran; LUX, Luxemburg; NDL, Netherlands; POL, Poland; RUS, Russia; USA, United States and ZAF, South Africa. **B**, Amino acid alignment of FgTPP1 for both protein haplotypes. The blue and brown squares represent predicted active sites and binding sites, respectively, between haplotype I and F. Annotations in gray, light violet, and orange lines represent the predicted signal peptide sequence (SP), the predicted trypsin domain, and the predicted chloroplast targeting signal, respectively. Isoleucine-4 (haplotype I) and phenylalanine-4 (haplotype F) indicate the difference between the two protein haplotypes. Functional secreted signals were predicted for both haplotypes using SignalP v6.0 (Teufel et al. 2022).

To evaluate the degree of sequence conservation of FgTPP1 in other fungal species, a BlastP search was conducted using the FgTPP1 protein sequence as the query. Homologs of FgTPP1 containing both a predicted secretion signal and a chloroplast transit peptide were identified not only in fungal species from the Fusarium genus, but also in fungal species belonging to different genera within the *Ascomycota* phylum (Figure 7 and Supplementary Tables S2 and S3). The FgTPP1 protein appears to be highly conserved (amino acid similarity values between 99 to 81%) in several plant pathogen species residing within the *Fusarium sambucinum* species complex including *F. culmorum*, *F. pseudograminearum*, *F. sporotrichioides*, *F. langsethiae*, *F. venenatum* and *F. poae* (Armer et al. 2024b; Waalwijk et al. 2018). In addition, the FgTPP1 protein is also conserved in other plant pathogenic Fusarium species (similarity values between 87 to 71%) such as *F. equiseti*, *F. fujikuroi*, *F. verticillioides*, *F. oxysporum* and *F. tricinctum* that belong to different Fusarium species complexes (Figure 7 and Supplementary Table S2). Some Fusarium species such as *F. sarcochroum*, *F. torreyae*, *F. tricinctum* and *F. avenaceum* possess two copies of the TPP1 gene (Supplementary Table S2). Finally, FgTPP1 is also conserved (similarity values between 64 to 52%) in fungal species from different genera within the *Ascomycota* phylum (Supplementary Table S3). Most of these species have been classified either as plant pathogenic (*Alternaria alternata* and *Verticillium dahliae*) or plant endophytic (*Colletotrichum tofieldiae* and *Alternaria rosae*) (Figure 2 and Supplementary Table S3). Multiple copies of TPP1 orthologs were also discovered in species residing in the Alternaria genus. While *A. panax* and *A. rosae* have 2 copies of *TPP1*, 3 copies were found in *A. alternata* and *A. burnsii*. The strong conservation of *TPP1* across a wide range of Ascomycete plant pathogens and endophytes suggests that TPP1 may play a conserved role(s) in plant colonization.

**Figure 7.**
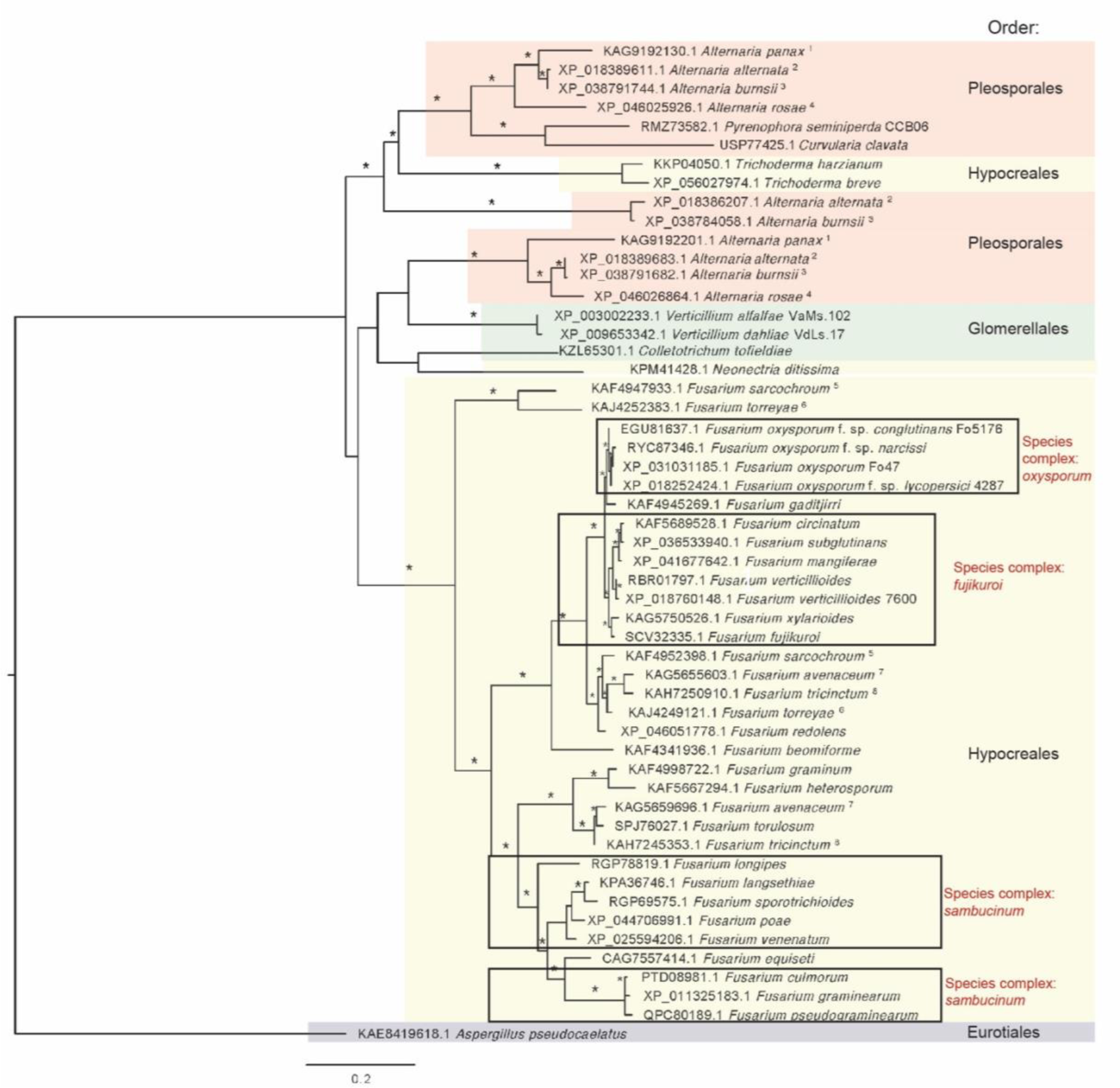
Phylogenetic tree of TPP1 protein homologs present in *Fusarium* species and other Ascomycete fungi. *Aspergillus pseudocaelatus* was used as an outgroup. Branches with asterisk (*) are supported by bootstrap value and/or posterior probability with minimum score 70/0.7. Multiple paralogs of TPP1 were present in single isolate of *Alternaria panax^1^*, *A. alternata^2^*, *A. burnsii^3^*, *A. rosae^4^*, *F. sarcochroum^5^*, *F. torreyae^6^*, *F. avenaceum^7^*, and *F. tricinctum^8^*.

## DISCUSSION

In this study, we examined the functional roles and immune suppressing properties of a putative effector protease from *F. graminearum*, FgTPP1, which was previously shown to be upregulated during early stages of wheat spike infection by *F. graminearum*, suggesting that is plays a role in fungal colonization and infection of the wheat spike. Consistent with this, the virulence of Δ*Fgtpp1* mutants was reduced when compared to the wild-type *F. graminearum* PH-1 strain using a bottom inoculation method. In this infection assay, *F. graminearum* colonizes the wheat rachis and vascular tissue, thereby inhibiting nutrient and water transport to the systemic, non-infected spikelets, resulting in bleaching of the wheat spike (Bai and Shaner 2004). During the bottom inoculation method, we observed fully bleached spikes and curved awns during the onset of spikelet bleaching (spike discoloration), which may be used as an additional criterion to categorize infection severity. Furthermore, these symptomatic characteristics allowed us to distinguish between fully bleached and partially bleached spikes, thereby affording us the opportunity to more consistently phenotype FHB disease progression. In contrast to the bottom inoculation method, the top inoculation approach consists of counting the number of diseased spikelets below the inoculation point and is often reliable when there are significant differences between mutant and wild type *F. graminearum* strains. However, distinguishing between an infected and uninfected wheat spikelet can be challenging using the top inoculation method, especially during the early stages of spikelet infection. Consequently, reduced virulence phenotypes are often difficult to reliably phenotype, especially when subtle. In our study, we did not observe a reduction in fungal virulence with the Δ*Fgtpp1-1* mutant when using the top inoculation method and the wheat cultivar ‘Bobwhite’. Our results presented here are also inconsistent with that of Menke (2011) who, in a non-peer reviewed study, showed that a Δ*Fgtpp1* mutant strain exhibited a subtle reduction in fungal virulence using the top inoculation method and the wheat cultivar ‘Norm’. Specifically, Menke (2011) reported that the number of diseased spikelets per infected spike with the wild-type *F. graminearum* PH-1 strain was 7.7 ± 0.3 while the number of disease spikelets per spike infected with the Δ*Fgtpp1* mutant was 7.2 ± 0.2, a 6% reduction in fungal colonization. Collectively, our experiments using different infection methods suggest that the bottom infection method is likely more sensitive to subtle virulence defects and could be used to further analyze *F. graminearum* mutants that do not show significant virulence defects using the top inoculation method.

Approximately seven predicted proteases, some of which are highly expressed during *in planta* fungal growth have been individually deleted in *F. graminearum*. However, only one *F. graminearum* mutant (FGSG_00192; FgPrb1) showed a significant reduction in fungal pathogenicity (Xu et al. 2020). This result may suggest that either candidate proteases are not directly involved in fungal pathogenicity, or fungal proteases may be functionally redundant, and, therefore, single deletion mutants are not sufficient to produce a strong reduction in fungal pathogenicity. For example, single gene deletions of two subtilisin-like protease genes, *FgSLP1* (FGSG_00806) and *FgSLP2* (FGSG_03315), did not significantly affect fungal virulence in wheat spikes when compared to wild-type *F. graminearum* PH-1 strain (Xu et al. 2020). However, simultaneous deletion of both *FgSLP1* and *FgSLP2* in resulted in a significant reduction in fungal virulence, indicating *FgSLP1* and *FgSLP2* are functionally redundant and complement each other during pathogenesis (Xiong et al. 2024). In addition to subtilisin-like proteases, *F. graminearum* expresses other putative proteases during *in planta* growth that attenuate plant defense responses, such as the serine carboxypeptidase FgSCP (FGSG_08454) (K. Liu et al. 2024). The FgSCP protease from *F. graminearum* suppresses cell death when transiently expressed in *N. benthamiana*, positively regulates the expression of DON biosynthesis, and plays a direct role in *F. graminearum* pathogenicity in wheat as well as maize (K. Liu et al. 2024). Similar to FgSCP1 and FgTPP1, the aspartic protease FolAsp, from *F. oxysporum f. sp. lycopersici*, was recently shown to inhibit hypersensitive response (HR)-like cell death as well as suppress PAMP-mediated ROS burst in *N. benthamiana* (Wang et al. 2023). Importantly, FolAsp is involved in *F. oxysporum* f. sp. *lycopersici* pathogenicity in tomato seedlings (Wang et al. 2023). The observation that *F. graminearum* expresses and secretes multiple fungal proteases that suppress plant immunity suggests deleting additional, functionally redundant proteases, in the Δ*Fgtpp1* mutant may further reduce fungal virulence.

Our finding that FgTPP1 localizes, in part, to the chloroplast stroma suggests *F. graminearum* may modulate chloroplast-mediated immune responses during fungal infection. It is worth noting that biosynthesis of the defense phytohormone salicylic acid (SA) occurs, in part, within chloroplasts and SA production is important for defense against biotrophic and hemibiotrophic pathogens (Ding and Ding, 2020; J. Liu et al. 2024). In addition to salicylic acid, chloroplasts also produce reactive oxygen species, which often function as signaling molecules to propagate immune responses against pathogens (Galvez-Valdivieso and Mullineaux, 2010; J. Liu et al. 2024). Considering the importance of SA production and ROS accumulation during plant-pathogen interactions, it stands to reason that downregulation of chloroplast functions could potentially interfere with production of defense signaling molecules (Kretschmer et al. 2020). Indeed, several filamentous fungal pathogens have evolved effector proteins that localize to chloroplasts to subvert chloroplast-derived defense responses (Figueroa et al. 2021; Littlejohn et al. 2021). For example, the wheat stripe rust pathogen *Puccinia striiformis* f. sp*. tritici* expresses and secretes the Pst_12806 effector inside host cells where it subsequently localizes to host chloroplasts and interacts with the chloroplast iron-sulfur protein, TaISP (Xu et al. 2019). The poplar rust fungus *Melampsora larici-populina* encodes three chloroplast-targeting proteins (CTP1, CTP2, and CTP3) that, similar to FgTPP1, contain chloroplast targeting sequences and localize to the stroma of chloroplasts in *N. benthamiana* (Petre et al. 2015, 2016). Although we hypothesize that the immune suppressing activities of FgTPP1 may be a result of its localization to chloroplasts, our data does not rule out the possibility that FgTPP1 may function in the nucleo-cytosol to suppress immunity. Hence, future work will focus on testing whether the subcellular localization of FgTPP1 is important for its immune-suppressing activities, identifying host proteins from wheat that interact with this candidate effector protease, and what effect such interactions have on facilitating *F. graminearum* infection and FHB disease development.

One of the reasons we have been researching effector proteases from *F. graminearum* is the potential to bioengineer novel recognition specificities of such proteases using decoy substrates. Many wheat and barley cultivars encode for the nucleotide-binding leucine-rich repeat (NLR) disease resistance gene *AvrPphB response 1* (*Pbr1*), which mediates recognition of the AvrPphB protease from *Pseudomonas syringae* (Carter et al. 2019; Jaiswal et al. 2023). AvrPphB is known to target receptor-like cytoplasmic kinases (RLCKs) in subfamily VII, which includes PBS1, from Arabidopsis (Shao et al. 2003; Zhang et al. 2010). We have previously shown that the AvrPphB cleavage site within PBS1 can be replaced with cleavage sequences for other pathogen-secreted proteases, including viral proteases, which then confers recognition and resistance to the pathogens expressing those proteases, including soybean mosaic virus and turnip mosaic virus (Helm et al. 2019; Kim et al. 2016). By extension, it may be feasible to introduce genetic-based resistance to *F. graminearum* by bioengineering a barley and wheat RLCK such that, when proteolytically cleaved by FgTPP1, activates PBR1-mediated immunity. While there are no reported NLR-based resistance genes against *F. graminearum*, we predict such NLR-mediated immune responses will be effective at conferring resistance to this fungal pathogen. For example, the tomato NLR immune receptor I2 was shown to confer recognition and resistance to *F. oxysporum* f. sp. *lycopersici* (Simons et al. 1998). A major goal for us going forward will be the identification of the preferred cleavage site sequence for FgTPP1, which would then enable us to generate decoy substrates that, when cleaved, would activate immune responses. If successful, the bioengineering of barley and wheat RLCK decoy proteins is thus expected to contribute to the development of improved crop protection strategies for *F. graminearum*. Importantly, we would expect such resistance to be quite durable, as *FgTPP1* is conserved among all sequenced *F. graminearum* genomes, which indicates that it likely has a central role in the *F. graminearum* life cycle. Furthermore, the conservation of *Fgtpp1* across fungal plant pathogens from the Ascomycetes phylum suggests that it may be possible to bioengineer resistance to fungal plant diseases of many different crops using the same approach.

## MATERIALS and METHODS

### Strain, media, and culture

*Fusarium graminearum* wild-type strain PH-1 (Cuomo et al. 2007) was used as the background strain to generate the Δ*Fgtpp1* mutant strains. The wild-type PH-1, PH-1-Δ*Fgtpp1* mutant strains, and the complementation strain PH-1-Δ*Fgtpp1-1*::*TPP1* were grown on synthetic nutrient agar (SNA; 0.1% KH_2_PO_4_, 0.1% KNO_3_, 0.1% MgSO_4_ x 7 H_2_O, 0.05% KCl, 0.02% glucose, 0.02% saccharose, and 2% agar) under constant ultraviolet and white light illumination. Conidiation was induced by adding 3 mL of TB3 liquid medium (0.3% yeast extract, 0.3% casamino acids, and 20% sucrose) to SNA plates containing 7-day old fungal mycelia. Conidia were harvested 2 days later in sterile water and stored at -80°C as described previously (Brown et al. 2011). *Escherichia coli* strain DH5ɑ was used for plasmid construction. *E. coli* transformants were selected on Luria-Bertani (LB) agar supplemented with either 25 µg/mL gentamicin or 100 µg/mL ampicillin. Defects in radial growth in the mutant and complemented strains compared to PH-1 were evaluated under different stress conditions as previously described (Darino et al. 2024). Briefly, 25mL of half-strength Potato Dextrose Agar (PDA) were mixed with different stress inducing agents such as membrane stresses (Congo Red [50 µg/mL], Calcofluor [100 µg/mL], and 0.02% Tergitol) and salt stress (1M NaCl) and poured into squares plates. The strains were also evaluated on SNA. Three 10-fold serial dilutions were prepared from a water stock containing 10^6^ fungal conidia/mL. Each spore dilution was spotted onto agar plates in 20µl droplets. Plates were incubated at room temperature (RT) under dark conditions for the entirety of the experiment. Photographs were taken at 2 days post inoculation (dpi). Defects in perithecia induction and formation between the PH-1-Δ*Fgtpp1-1* mutant strain and PH-1 were evaluated on carrot agar medium following the protocol described by Cavinder et al. (2012). Photographs were taken 9 days after inoculation of the fungal strains on plates containing carrot agar. The experiment was repeated three independent times.

### Plant growth conditions

Seeds of *Nicotiana benthamiana* were grown in pots containing Berger Seed and Propagation Mix supplemented with Osmocote slow-release fertilizer (14-14-14) and maintained in a growth chamber with a 16:8 h photoperiod (light:dark) at 24°C in the light and 20°C in the dark, with average light intensities at plant height of 120 µmols/m^2^/s.

Spring wheat cultivar ‘Bobwhite’ seedlings were grown in pots containing Rothamsted mix soils with 50% the standard fertilizer rate and maintained in a controlled environment room with a 16:8 h photoperiod (light:dark) under white-based LEDs with far-red addition (HelioSPEC R40F Flex, Sweden) with average light intensities of 300 µmols/m^2^/s and 65% relative humidity. Temperatures during light/dark conditions were 20°C and 18°C, respectively (Darino et al. 2024).

### Prediction of the FgTPP1 protein structure using AlphaFold2

The protein structure of FgTPP1 (without the predicted signal peptide sequence) was predicted using the ColabFold v1.5.2: AlphaFold2 using MMseqs2 with default parameters as previously described (Mirdita et al. 2022; Rogers et al. 2024). The predicted protein structure with the highest confidence (pLDDT) score as determined by AlphaFold2 (i.e. ranked_0.pdb) was captured and visualized using ChimeraX-1.5 (Meng et al. 2023).

### Yeast secretion trap assay for testing signal peptide function

Secretion of different *F. graminearum* proteins was tested in yeast using the yeast secretion trap assay as previously described by Zhou and colleagues (2020) with slight modifications. Briefly, secretion signals from FgTPP1 and FgOSP24 were predicted using SignalP v6.0 (Teufel et al. 2022). Full length coding sequences of *FgTPP1*, *FgOSP24*, and *FgTPP1* without the predicted secretion signal (*FgTPP1^ΔSP^*) were PCR-amplified using primer combinations P1 / P2, P3 / P4, and P5 / P2, respectively, using cDNA generated from wheat floral tissue infected with *F. graminearum* wild type strain PH-1 as the template. The resulting PCR products were cloned into the pDONR207 plasmid using a Gateway cloning approach (Invitrogen). Donor clones containing pDONR207-*FgTPP1*, pDONR207-*FgOSP24*, and pDONR207-*FgTPP1^ΔSP^* were recombined with the destination vector pGAD-GW-SUC2^22-511^ using the Gateway cloning system (Invitrogen) to generate the following constructs: pGAD-*FgTPP1*:SUC2^22–511^, pGAD-*FgOSP24*:SUC2^22–511^, and pGAD-*FgTPP1^ΔSP^*:SUC2^22–511^. Each construct was transformed into a yeast sucrose invertase (*suc2*) mutant using a yeast transformation kit following the manufacturer’s instructions (Yeast Transformation Kit, Sigma-Aldrich). The *suc2* mutant is unable to secrete the SUC2 protein and thus cannot grow on media containing sucrose (Suc) as the sole carbon source. Transformants were plated on yeast synthetic dropout media (SD) lacking tryptophan and leucine (-TL) supplemented with 2% glucose (Glu). The lack of tryptophan and leucine is to select positive transformants of the yeast *suc2* mutant containing the different constructs. Plates were incubated at 30°C for 3 days. Yeast transformants were verified by PCR amplification to determine the presence of the fusion proteins using the primer combination P6 and P7. Positive clones were inoculated into liquid SD-TL 2% Glu media and incubated at 30°C overnight with slight agitation (180 rpm). Following overnight incubation, liquid cultures were adjusted to an optical density (OD) at 600 nm (OD_600_) of 1.0 and centrifuged at 4000 rpm for 2 minutes to remove the growth media. Yeast pellets were resuspended in sterile water and three 10-fold serial dilutions for each transformant were prepared. Five microliters of each dilution was spotted onto plates containing SD-TL supplemented with either 2% Glu or 2% Suc. Plates were incubated at 30°C for 4 days. Empty vector pGAD-GW-SUC2^22-511^ was used as negative control. The experiment was repeated twice.

### Deletion of *tpp1* and complementation in *F. graminearum*

The **‘**split-marker’ approach utilizing two overlapping DNA fragments was used to delete the *FgTPP1* gene from the *F. graminearum* PH-1 strain (Catlett et al. 2003; King et al. 2017b). Briefly, two plasmids, pEK01 and pEK02, were designed to delete *FgTPP1*. The vector pEK01 contained a 1,000 base pair fragment upstream of the start codon of the *FgTPP1* gene (P_TPP1_) followed by a partial sequence of the hygromycin B resistance cassette (Hyg_1-761_). P_TPP1_ was PCR-amplified from *F. graminearum* PH-1 genomic DNA using primers P8 and P9, whilst the Hyg_1-761_ fragment was PCR-amplified from the pHyg1.4 vector using primers P10 and P11 (Urban et al. 2003). The vector pEK02 contained a fragment of the hygromycin B resistance cassette (Hyg_296-1027_) followed by a 1,000 base pair fragment of the terminator region of *FgTPP1* downstream of the stop codon (T_TPP1_). Primers P12 and P13 were used to PCR amplify T_TPP1_ from PH-1 genomic DNA whilst primers P14 and P15 were used to PCR amplify the fragment Hyg_296-1027_ from the pHyg1.4 vector. Gibson assembly was used to ligate the PCR fragments (P_TPP1_ with Hyg_1-761_ and Hyg_296-1027_ withT_TPP1_) into the pGEM-T Easy vector to generate plasmids pEK01 and pEK02, respectively. The two plasmids share a 466 bp overlapping region of the Hyg cassette to allow recombination during the transformation step.

To proceed with the transformation step, PCR products containing the P_TPP1_-Hyg_1-76_ and Hyg_296-1027_-T_TPP1_ were PCR-amplified using HotStart polymerase and primer combinations P8 / P11 and P14 / P13, respectively. Then, 5 µL aliquots of 2 µg/mL from each PCR product were combined and used to transform 1 × 10^8^ protoplasts generated from *F. graminearum* PH-1 according to the protocols described by Hohn and Desjardins (1992) and King et al. (2017b). Transformants were selected on regeneration media (0.2% yeast extract, 0.2% casein-hydrolysate (N-Z-Amine A), 0.7% agarose and 0.8 M sucrose) containing 75 µg/mL hygromycin. Well-spaced transformants were selected using sterile wooden toothpicks and transferred to 6-well plates containing SNA with 75 µg/mL hygromycin. To extract genomic DNA, transformants were grown on YPD broth containing 75 µg/mL Hyg B and DNA was extracted according to the method described by Rudd et al. (2010). PCR diagnostic tests were performed to confirm the correct insertion of the Hyg cassette into the TPP1 locus (primer combinations: P16 / P17 and P18 / P19) and to test for *FgTPP1* coding sequence replacement (primer combination P20 / P21) (King et al. 2017b).

The Δ*Fgtpp1-1* mutant was complemented with the coding sequence of *FgTPP1* at the TSI locus 1 (Darino et al. 2024). A DNA fragment containing the *FgTPP1* coding sequence, including the promoter and terminator regions of *FgTPP1* (P*_tpp11_*-*TPP1*-T*_tpp1_*) was PCR-amplified from genomic DNA of PH-1 using primer combination P22 and P23. The PCR product was cloned into the Fg vector as previously described (Darino et al. 2024). The Fg vector contains a fragment of the geneticin resistance cassette followed by the right border (RB) of the TSI locus 1 (*geneticin*_1-664_-RB). Between the RB and the partial sequence of the geneticin cassette, there is a cloning site adapted to the Golden Gate cloning approach (Engler et al. 2009). Integration into the TSI locus 1 was accomplished following an adaptation of the split-marker approach (Darino et al. 2024). Briefly, a PCR product containing the partial fragment of the geneticin resistance cassette followed by the P*_tpp1_*-*TPP1*-T*_tpp1_* region and the RB of the TSI locus 1 (*geneticin*_1-664_-P*_tpp1_*-*TPP1*-T*_tpp1_*-RB) was PCR-amplified from the Fg vector using primer combination P24 and P25. A second PCR product containing the left border (LB) of the TSI locus 1 followed by the split fragment of the geneticin resistance cassette (*geneticin*_795-128_) was amplified from the pJET-LB-geneticin vector using primer combination P26 and P27. Both PCR products possess an overlapping region of 536 bp to allow homologous recombination. The PCR products were adjusted to a final concentration of 2 µg/µL. Finally, 5 µL of each PCR product was aliquoted and mixed. The mixture of PCR products was used to transform *F. graminearum* protoplasts prepared from the PH-1-Δ*Fgtpp1-1* mutant following the protocol described above. Transformants were selected on SNA plates containing 75 µg/mL of geneticin. To test for single insertion of the expression cassette into the TSI locus 1, a diagnostic PCR spanning from outside the recombination region was performed using primer combination P28 and P29. A transformant with an amplicon of 6,123 bp was selected as the positive transformant.

### FHB virulence assays

The susceptible spring wheat cultivar ‘Bobwhite’, sourced from the International Maize and Wheat Improvement Centre (CIMMYT, Mexico), was inoculated at anthesis following the top inoculation approach as previously described by Wood et al. (2020) with slight modifications. Briefly, two spikelets per wheat spike were inoculated, using 4 to 7 plants per fungal strain and only the first spike per plant. The 14^th^ and 15^th^ spikelets, counting up from the bottom, were point-inoculated with 5µL of 1x10^5^ conidia/mL of either PH-1-Δ*Fgtpp1-1* or wild-type *F. graminearum* PH-1. As a control, wheat cv. ‘Bobwhite’ plants were inoculated with sterile, deionized water. Inoculated wheat plants were incubated under high humidity conditions (above 80%) for 48 hours, with an initial 24 hours in the dark. Fusarium Head Blight disease was scored by counting the number of infected spikelets below the point of inoculation every 2 days until 12 days post-inoculation. Area under the disease progress curve (AUDPC) analysis was used to quantify statistical differences between the mutant and PH-1 according to Schandry (2017). The AUDPC values were calculated for each spike infected with PH-1-Δ*Fgtpp1-1* or PH-1. Then, AUDPC average values for PH-1-Δ*Fgtpp1-1* and PH-1 were calculated and significant differences between PH-1-Δ*Fgtpp1-1* mutant and PH-1 were calculated using a t test with p< 0.05.

A bottom-inoculation approach was also performed to assess the virulence of the PH-1-Δ*Fgtpp1* mutant and PH-1-Δ*Fgtpp1-1*::*TPP1* complementation strains compared to wild-type PH-1. This approach is an adaptation of a previously described protocol (Seong et al. 2005). The 3^rd^ and 4^th^ full sized spikelets from the bottom of the wheat spike were inoculated, one spike per plant, and 5 to 20 plants per fungal strain. The two spikelets at the base were point-inoculated with 5µL of 5x10^4^ conidia/mL of either PH-1-Δ*Fgtpp1-1,* PH-1-Δ*Fgtpp1-3*, PH-1-Δ*Fgtpp1-1*::*TPP1*, or PH-1. Inoculated plants were incubated as described above. Between 10- and 11-days post-inoculation and depending upon the disease severity per infection batch, wheat spikes were classified as either completely bleached (wherein all the spikelets were found to be bleach and the awns spread out) or partially bleached (wherein some but not all the spikelets were bleached, and most awns remained upright). A goodness of fit test was performed to test if the observed frequencies of fully bleached / partially bleached spikes for each fungal strain were the same or different. The experiment was repeated at least four independent times. Images were captured using Olympus OM-D E-M10 fitted with a M.Zuiko 30mm F3.5 macro lens.

### DON mycotoxin measurements

DON mycotoxin concentrations within wheat spikes infected either with the *ΔFgtpp1-1* mutant strain or PH-1 were assessed using the Deoxynivalenol (DON) Plate Kit (Beacon Analytical Systems, Inc., US). Briefly, top inoculated wheat spikes infected with wild-type *F. graminearum* PH-1 or the PH-1-*ΔFgtpp1-1* mutant strain were collected at 18 days post-inoculation and frozen in liquid nitrogen. Three independent spikes per strain were individually grounded to a fine powder. Nine volumes of sterile water were added to the ground tissue. Then, the solution was mixed by vortexing and the supernatant was collected after centrifugation at maximum speed (12,000 rpm). The supernatant was ten-fold diluted and 50 µl of this diluted supernatant was mixed with 50 µl of the enzyme conjugate following the protocol described by the manufacturer. Three independent mock-inoculated wheat spikes were also included as a negative control. A calibration curve using different DON concentrations was used to determine the DON parts per million (ppm) for each wheat spike. A t test analysis was performed to calculate significant differences between the PH-1-*ΔFgtpp1-1* mutant and PH-1 strains (p value < 0.05).

### Generation of plant expression constructs

The RPS5^D266E^:myc and RbcS-TP:mCherry constructs have been described previously (Helm et al. 2022; Qi et al. 2012).

To generate the FgTPP1:GFP and FgTPP1:mCherry constructs, we used the Golden Gate protocol as previously described by Lampropoulos et al. (2013). Briefly, the *FgTPP1* open reading frame, without the predicted signal peptide and the stop codon, was PCR-amplified using primer combination P30 and P31 using cDNA isolated from wheat floral tissue infected with PH-1 as the template. The resulting PCR fragments were purified and combined in a single Golden Gate reaction with the destination vector pGGZ001 for *in planta* expression. The pGGZ001 plasmid contains a 35S promoter, Ω-element (protein expression enhancer), UBQ10 terminator, HygR resistant cassette and either mCherry or GFP as C-terminus tags. The resulting constructs were sequence confirmed and subsequently designated as FgTPP1:GFP (pGGZ001-35s-Ω-element-*FgTPP1*:GFP-UBQ10t-HygR) and FgTPP1:mCherry (pGGZ001-35s-Ω-element-*FgTPP1*:mCherry-UBQ10t-HygR). A free GFP construct (pGGZ001-35s-Ω-element-*GFP*-UBQ10t-HygR) was also generated using the aforementioned Golden Gate protocol.

To generate the FgTPP1:sYFP construct, the *FgTPP1* open reading frame (without the signal peptide sequence) was synthesized by TWIST Biosciences with codon optimization for expression in *N. benthamiana* and cloned into pTWIST-Kan by the service provider. This ORF was then PCR-amplified with *attB*-containing primers. The resulting PCR fragments were purified and recombined into the Gateway entry vector, pBSDONR(P1-P4) (Qi et al. 2012) using BP Clonase II (Invitrogen) and the resulting construct was designated pBSDONR(P1-P4):*FgTPP1*. The pBSDONR(P1-P4):*FgTPP1* plasmid was mixed with the pBSDONR(P4r-P2):*sYFP* (Qi et al. 2012) plasmid and the Gateway-compatible expression vector pTA7001 (Vinatzer et al. 2006). Plasmids were recombined by the addition of LR Clonase II (Invitrogen). The resulting construct was designated pTA7001:*FgTPP1*:sYFP. All constructs were sequence-verified for proper sequence and reading frame.

### *Agrobacterium*-mediated transient expression in *Nicotiana benthamiana*

Transient protein expression assays were performed as previously described with minor modifications (Helm et al. 2022). Briefly, the constructs described above were mobilized into *Agrobacterium tumefaciens* GV3101 (pMP90) and grown on Luria-Bertani (LB) media plates supplemented with 25 µg/ml of gentamicin sulfate and 50 µg/ml of kanamycin for two days at 30°C. Cultures were prepared in liquid LB media supplemented with the appropriate antibiotics and were shaken overnight at 30°C on an orbital shaker. Following overnight incubation, cells were pelleted by centrifuging at 3000rpm for 3 minutes at room temperature. For confocal microscopy and immunoblot analyses, bacterial pellets were resuspended in 10 mM MgCl_2_, adjusted, and mixed to an OD_600_ of 0.3 for each strain, and incubated with 100 µM of acetosyringone for 3-4 hours at room temperature. For the suppression of HR-like cell death assay, bacterial pellets of FgTPP1:sYFP and RPS5^D266E^:myc were resuspended in 10 mM MgCl_2_, adjusted to an OD_600_ of 0.3 and 0.15, respectively, and incubated with 150 µM of acetosyringone for 3 hours at room temperature. All bacterial suspensions were infiltrated into the abaxial side of *N. benthamiana* leaves using a needless syringe. For pTA7001-based constructs, which are dexamethasone-inducible, gene expression was induced approximately 48 hours post-agroinfiltration by spraying leaves with 50 µM dexamethasone.

### Confocal microscopy

Confocal microscopy of *N. benthamiana* epidermal cells was performed twenty-four hours post-agroinfiltration using a Zeiss LSM880 Axio Examiner upright confocal microscope with a Plan Apochromat 20x/0.8 objective. Fluorescence from the GFP-tagged protein fusions was excited using a 488-nm argon laser and detected between 525-nm and 550-nm. Fluorescence from the mCherry-tagged constructs was excited with a 561-nm helium-neon laser and detected between 565-nm and 669-nm. All confocal micrographs shown are of single optical sections and processed using the Zeiss Zen Blue Lite program (Carl Zeiss Microscopy).

### Immunoblot analyses

*N. benthamiana* leaves transiently expressing the epitope-tagged constructs were harvested 24 or 48 hours following agroinfiltration, flash frozen in liquid nitrogen, and total protein extracted as previously described by Helm et al. (2022) for immunoblot analyses. Ten microliters of isolated total protein were separated on a 4-20% Tris-glycine stain free polyacrylamide gel (Bio-Rad) at 170 V for one hour in 1X Tris/glycine/SDS running buffer. Total proteins were transferred to nitrocellulose membranes (GE Water and Process Technologies). Membranes were washed with 1X Tris-buffered saline (50 mM Tris-HCl, 150 mM NaCl, pH 7.5) solution supplemented with 0.1% Tween20 (TBST) and blocked in 5% skim milk (w/v) (Becton, Dickinson & Company) for at least 1 hour at room temperature. Proteins were detected with horseradish peroxidase (HRP)-conjugated anti-GFP (1:5,000; Miltenyi Biotec) antibodies for one hour at room temperature. Membranes were washed three times with 1X TBST solution and subsequently incubated with Clarity Western ECL (Bio-Rad) substrate solution for 5 minutes. Imaging of immunoblots was performed with an ImageQuant 500 CCD imaging system.

### Chloroplasts isolation

Chloroplast isolation was performed as previously described with slight modifications (Petre et al. 2016). *N. benthamiana* leaves transiently expressing the indicated epitope-tagged constructs were harvested 24 hours following agroinfiltration, cut into 1 mm^2^ pieces, and incubated in 20 mL of cold isolation buffer (IB) (400 mM sorbitol, 50 mM of HEPES KOH 8M, 2 mM EDTA, and 1 mM MgCl_2_) for 15 minutes on ice. Leaf pieces were homogenized with a PT 1200 C Polytron homogenizer (3 × 5 s full speed), and the lysate was immediately filtered through doubled Miracloth to remove cell debris. The filtrate was collected in 50 mL conical tubes and centrifuged at 3000 rpm for 10 minutes at 4°C. The pellet containing the organelles was resuspended in 400 µl of ice-cold isolation buffer, and chloroplasts were isolated by centrifugation on a Percoll gradient (1 mL of 80% v/v Percoll / IB and 3 mL of 40% v/v Percoll/IB in a 10 mL conical tube) at 4500 rpm for 10 minutes at 4°C. Intact chloroplasts were collected from the bottom layer and subsequently used for laser scanning confocal microscopy.

### MAP kinase (MAPK) activity assay

Suppression of chitin-induced MAPK activation in *N. benthamiana* was performed as previously described with slight modifications (Jaiswal et al. 2022). Briefly, *N. benthamiana* leaves transiently expressing either free GFP or GFP-tagged TPP1 (FgTPP1:GFP) were infiltrated with either nuclease free water (mock treatment) or chitin hexamer dissolved in nuclease free water (5 μg/mL; Accurate Chemicals & Scientific Corporation). Leaf discs (10 mm diameter) were harvested and frozen in liquid nitrogen at 0-, 5-, and 10-minutes following mock or chitin infiltration. Total protein was extracted as described above. Proteins were detected using HRP-conjugated anti-GFP (1:5,000; Miltenyi Biotec), anti-plant actin (1:5000; Abbkine), or anti phospho-P42/44 MAPK (1:5000; Cell Signaling Technology), which detects phosphorylated MPK3 and MPK6. Membranes were washed three times with 1X TBST solution and subsequently incubated with Clarity Western ECL (Bio-Rad) substrate solution for 5 minutes. Imaging of immunoblots was performed with an ImageQuant 500 CCD imaging system. Experiments were performed at least three independent times with similar results.

### Luminol-based reactive oxygen species (ROS) assay in *N. benthamiana*

Suppression of ROS production in *N. benthamiana* was performed as previously described using a luminol-based chemiluminescence assay (Jaiswal et al. 2022; Rogers et al. 2024). *N. benthamiana* leaf discs (5 mm diameter) transiently expressing either free GFP or FgTPP1:GFP were harvested two days post-agroinfiltration using a cork borer, gently washed three times in nuclease free water, and incubated overnight in sterile water in a 96-well OptiPlate^TM^ microplate (Perkin Elmer). The following day, the deionized water was replaced with chitin elicitation solution (luminol [30 μg/mL], horseradish peroxidase [20 μg/mL], chitin hexamer [5 μg/mL] (Accurate Chemicals & Scientific Corporation), and nuclease-free water) and ROS production was monitored by chemiluminescence for 40 minutes in a microplate reader (Tecan Infinite M200 Pro). Experiments were performed at least three independent times with similar results.

### Cell death suppression assay

Forty-eight hours following agroinfiltration with either empty vector (e.v.), FgTPP1:sYFP, or RPS5^D266E^:myc, *N. benthamiana* leaves were sprayed with 50 μM dexamethasone supplemented with 0.02% Tween20 (to induce expression of both RPS5^D266E^:myc and FgTPP1:sYFP). At 16 hours post-transgene induction, *N. benthamiana* leaves were assessed for cell death and photographed under white and ultraviolet (UV) light. Experiments were performed at least three independent times with similar results.

### TPP1 sequence analysis from *F. graminearum* global populations and phylogenetic analyses

Haplotype analysis of FgTPP1 nucleotide sequences from *F. graminearum* isolates was conducted according to Darma (2023). In brief, *FgTPP1* nucleotide sequences from *F. graminearum* isolates available in the NCBI database were extracted using BLAST+ v2.9.0+ (Camacho et al. 2009) and parsed into a FASTA fasta file using ‘BLASTtoGFF_50percent.py’ (https://github.com/megancamilla). *FgTPP1* sequences were aligned using Geneious 10.2.3 alignment with the Blosum62 cost matrix. Introns from those sequences were trimmed to generate coding sequences of *FgTPP1* from global populations. Allele networks from *FgTPP1* coding sequences were generated with POPart v1.7 (Leigh and Bryant, 2015) using minimum spanning network setting.

TPP1 protein sequences from *Fusarium* sp. and other fungi were extracted with BlastP. Multiple hits representing TPP1 homologs from different *Fusarium* species complexes were selected for further analysis. In addition, hits from other fungi, excluding *Fusarium* species, predicted as trypsin or serine protease with the minimum max score 243, were extracted for further analysis. TPP1 protein sequences from *Fusarium* species and other fungi were aligned using Geneious 10.2.3 alignment as described previously. IQ-Tree 2.2.2.6 (Kalyaanamoorthy et al. 2017; Nguyen et al. 2015) was used to identify the best amino acid substitution model from the TPP1 amino acid sequence alignment and was used to generate a maximum likelihood tree using 10,000 bootstrap replicates. In addition, a Bayesian inference tree was also generated to support the maximum likelihood tree using MrBayes 3.2.6 (Huelsenbeck and Ronquist, 2001) with the following settings: rate matrix (fixed) WAG, rate variation gamma, gamma categories 4, chain length 1,000,000, heated chains 4, heated chain temperature 0.2, subsampling frequency 10,000, burn-in length 100,000, and random seed 16,291. Finally, Figtree 1.4.4 (Rambaut 2018) was used to refine the appearance of the phylogenetic tree.

## ACKNOWLEDGEMENTS

We thank Dr. Tesfaye Mengiste (Purdue University) for access to the microplate reader for the ROS suppression assays. We would also like to thank Dr. Terri Cameron for technical assistance, and the Purdue University Imaging Facility for access to the Zeiss LSM880 Axio Examiner upright confocal microscope. The authors would also like to thank Dr. Fiona Doohan (University of Dublin, Ireland) for providing the pGAD-SUC2^22-511^ plasmid and the *suc2* yeast mutant, Silvia Melina Velasquez for her technical assistance with the violin plots, Fiona Gilzean and the facilities team at Rothamsted Research for preparing and maintaining the various growth rooms used in this study whilst major refurbishment of the containment facilities were in progress. All experiments involving *F. graminearum* strain PH-1 and isogenic transformants were conducted in biological containment facilities under UK Defra license number 101948/198285. All opinions expressed in this paper are the author’s and do not necessarily reflect the policies and views of USDA. Mention of trade names or commercial products in this publication is solely for the purpose of providing specific information and does not imply recommendation or endorsement by the U.S. Department of Agriculture. USDA is an equal opportunity provider and employer.

## SUPPLEMENTARY FIGURES

**Supplementary Figure S1.**
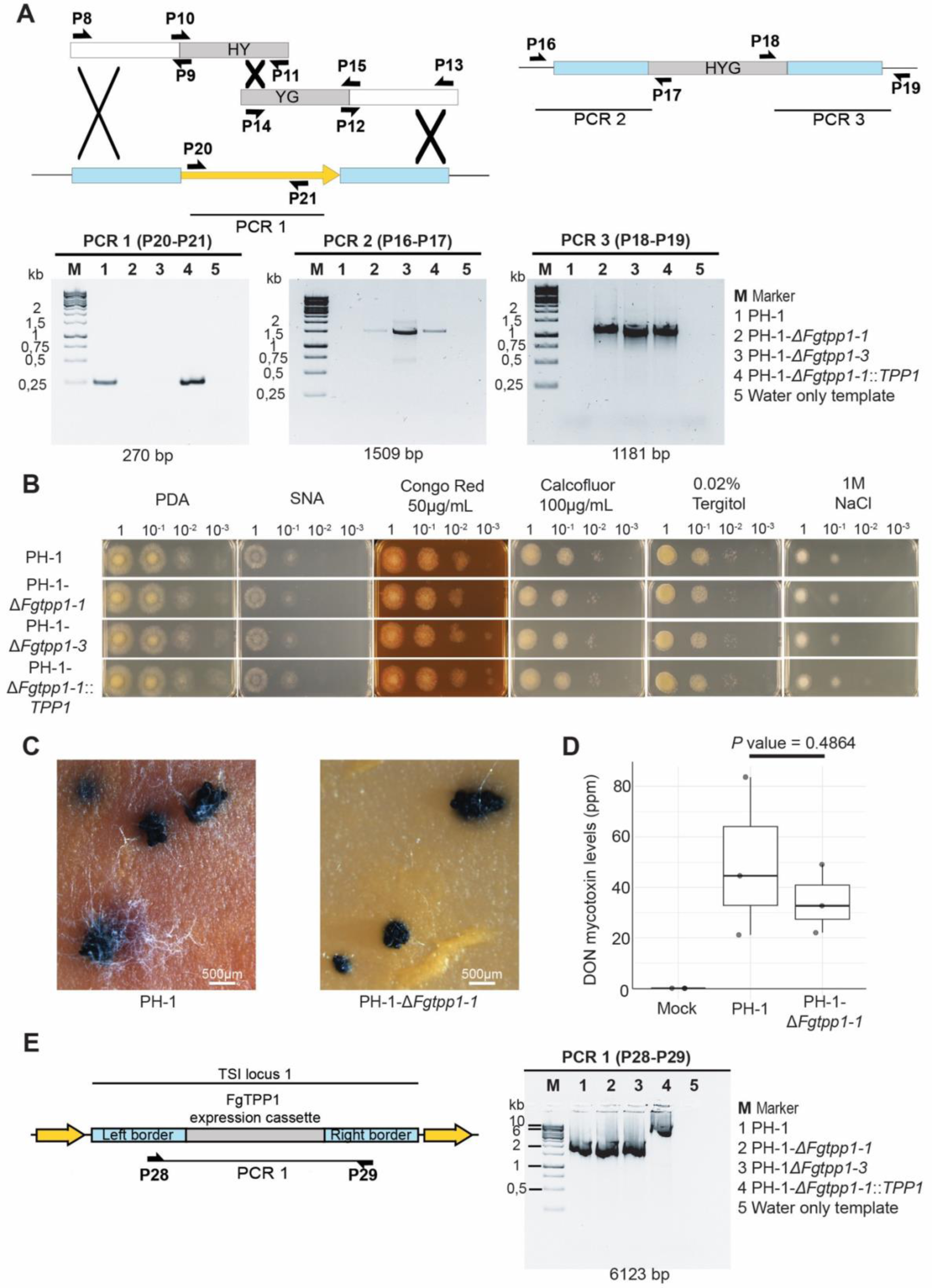
*In vitro* stress tests and DON quantification for wild-type *F. graminearum* PH-1 strain, the *ΔFgtpp1* mutant strains (PH-1-*ΔFgtpp1-1* and PH-1-*ΔFgtpp1-3*, and the complementation strain (PH-1-*ΔFgtpp1-1*::*TPP1*). **A,** Multiple PCR amplifications using various primer combinations were performed to select two independent fungal strains in which the *FgTPP1* gene had been successfully deleted. PCR 1 (primer combination P20 and P21) evaluated the *FgTPP1* coding sequence deletion. PCR2 (primer combination P16 and P17) and PCR3 (primer combination P18 and P19) confirmed correct insertion of the Hygromycin cassette into the *FgTPP1* locus. **B,** Stress tests for wild-type *F. graminearum* PH-1 strain, both *ΔFgtpp1* mutants, and the *ΔFgtpp1* complementation strain. The mutant and the complementation strains showed growth rates and morphology indistinguishable from that of the wild-type *F. graminearum* PH-1 strain for all the conditions evaluated. Images were taken 2 days post-plating on plates. PDA, Potato dextrose agar; SNA, Synthetic nutrient agar; Membrane stresses: Congo Red, Calcofluor, and Tergitol; and salt stress (NaCl). **C**, No obvious defects were observed in perithecia induction and formation between the PH-1-*ΔFgtpp1-1* mutant and the wild-type *F. graminearum* PH-1 strain. Perithecia images were taken 9 days after perithecia induction on carrot agar medium. **D**, No significant differences in DON mycotoxin production were observed between the PH-1-*ΔFgtpp1-1* mutant and wild-type *F. graminearum* PH-1 strain. DON quantification was performed using wheat spikes infected with either PH-1-*ΔFgtpp1-1* mutant or wild-type *F. graminearum* PH-1. Mock inoculated spikes were used as a negative control. Three independent wheat spikes were evaluated for each fungal strain and mock control (t test, p value > 0.05). **E**, Diagnostic PCR for the Δ*Fgtpp1* complementation strain (PH-1-ΔFg*tpp1-1*::*TPP1)*. A PCR was performed using primer combination P28 -P29 to evaluate correct insertion of the TPP1 expression cassette containing the TPP1 coding sequence and its native promoter and terminator in the TSI locus 1. PH-1 and the *ΔFgtpp1* mutant strains showed a 2100 base pair amplicon whilst the complementation strain showed an amplicon of 6123 base pair, which indicated insertion of the TPP1 cassette in the TSI locus 1.

**Supplementary Figure S2.**
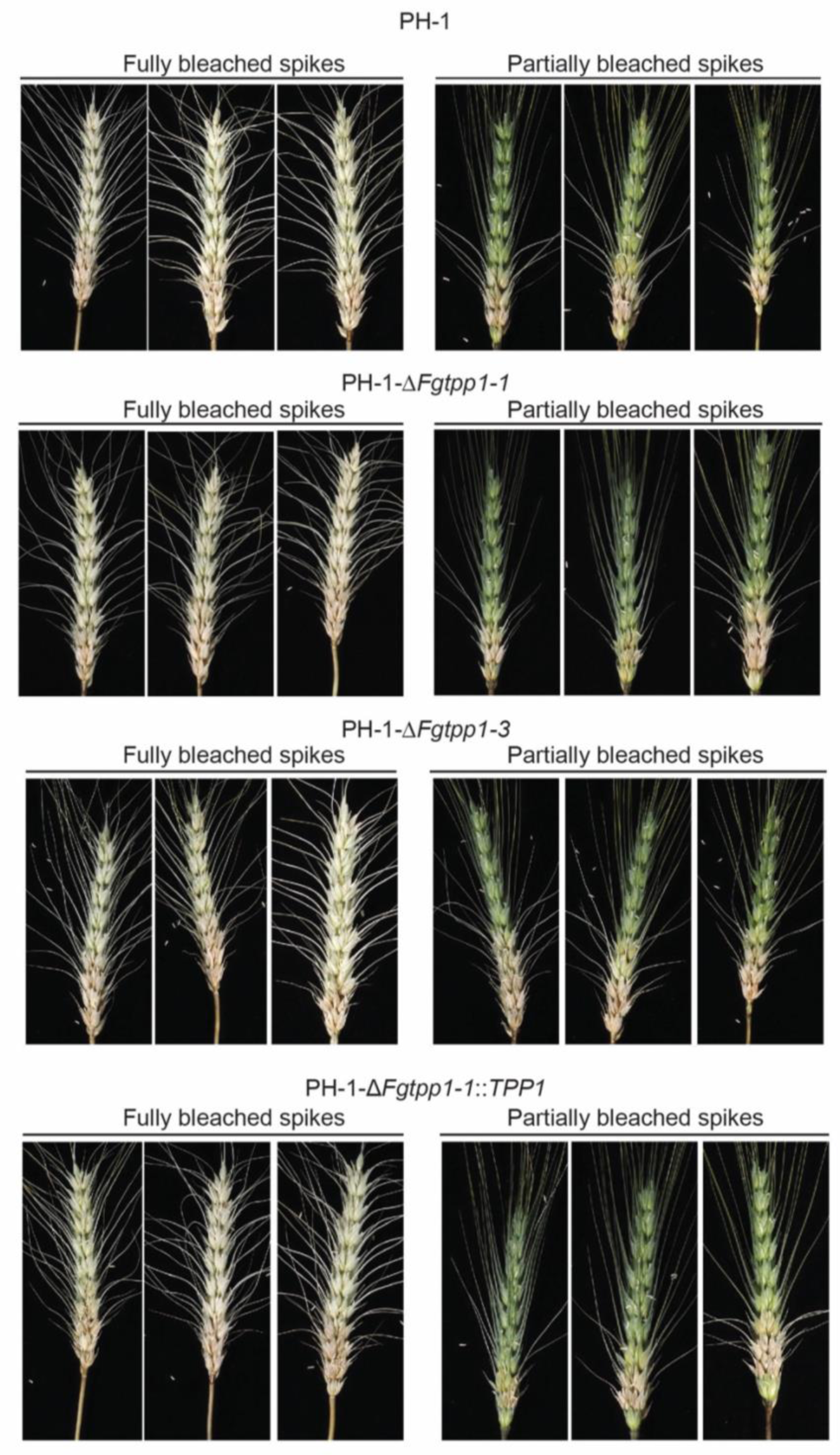
Representative images of infected wheat spikes using the bottom inoculation method photographed 10 days post-infection. Images belong to a single replicate where wild-type *F. graminearum* PH-1, the *ΔFgtpp1* mutant strains (PH-1-*ΔFgtpp1-1* and PH-1-*ΔFgtpp1-3*), and the complementation strain (PH-1-*ΔFgtpp1-1*::*TPP1*) were evaluated.

**Supplementary Table S1.** Accession numbers for the different *F. graminearum* strains included in the FgTPP1 nucleotide and protein haplotype analyses.

| Strain name | Country of origin | Year isolated | NCBI project accession number |
| --- | --- | --- | --- |
| PH-1 | USA | 2000 | PRJNA13839 |
| KSU23473 | USA | 2000 | PRJNA989623 |
| KSU23522 | USA | 2000 | PRJNA989623 |
| KSU23468 | USA | 1999 | PRJNA989623 |
| KSU23389 | USA | 2013 | PRJNA989623 |
| NRRL 29169 | USA | 1990 | PRJNA527821 |
| 233423 | USA | 1995 | PRJNA274567 |
| DAOM 241165 | Canada | 2004 | PRJNA274567 |
| CML3066 | Brazil | 2009 | PRJEB12819 |
| CS3005 | Australia | 2000 | PRJNA235346 |
| N4_2 | Germany | 2016 | PRJNA677929 |
| ITA_285 | Italy | 1993 | PRJNA677929 |
| ITA_1601 | Italy | 2001 | PRJNA677929 |
| ITA_202 | Italy | 1993 | PRJNA677929 |
| CBS 119799 | South Africa | 1987 | PRJNA677929 |
| CBS 128539 | Belgium | 2007 | PRJNA677929 |
| CBS 110263 | Iran | 1968 | PRJNA677929 |
| 114-2 | Argentina | 2012 | PRJNA677929 |
| C10a | France | 2002 | PRJNA677929 |
| 87-7 | Argentina | 2011 | PRJNA677929 |
| S5A | France | 2008 | PRJNA677929 |
| 14C1 | Netherlands | 2000 | PRJNA677929 |
| 0343 | Poland | 2003 | PRJNA677929 |
| 58918 | Russia | 2015 | PRJNA677929 |
| S17-12 | Poland | 2017 | PRJNA677929 |
| Fg820 | Netherlands | 1999 | PRJNA677929 |
| 237 | Luxembourg | 2007 | PRJNA677929 |
| FG_08 | Germany | 1987 | PRJNA677929 |

**Supplementary Table S2.** FgTPP1 protein orthologs in the *Fusarium* genus.

| Accession number<sup>1</sup> | Fungal species | Fusarium species complex<sup>2</sup> | Lifestyle | Query Coverage | Identity (%) |
| --- | --- | --- | --- | --- | --- |
| PTD08981.1 | *F. culmorum* | *Sambucinum* | Plant pathogen | 100% | 99.21 |
| QPC80189.1 | *F. pseudograminearum* | *Sambucinum* | Plant pathogen | 100% | 98.41 |
| CAG7557414.1 | *F. equiseti* | *Incarnatum - equiseti* | Plant pathogen | 90% | 87.72 |
| XP_025594206.1 | *F. venenatum* | *Sambucinum* | Plant pathogen | 91% | 86.15 |
| RGP69575.1 | *F. sporotrichioides* | *Sambucinum* | Plant pathogen | 100% | 86.11 |
| KPA36746.1 | *F. langsethiae* | *Sambucinum* | Plant pathogen | 100% | 86.11 |
| XP_044706991.1 | *F. poae* | *Sambucinum* | Plant pathogen | 100% | 82.14 |
| RGP78819.1 | *F. longipes* | *Sambucinum* | Plant endophytic | 99% | 81.27 |
| XP_046051778.1 | *F. redolens* | *Redolens* | Plant pathogen | 91% | 78.35 |
| KAH7250910.1 | *F. tricinctum* | *Tricinctum* | Plant pathogen | 91% | 76.62 |
| KAJ4249121.1 | *F. torreyae* | No complex assigned | Plant pathogen | 100% | 76.59 |
| KAF4952398.1 | *F. sarcochroum* | *Lateritium* | Not known | 91% | 76.19 |
| KAF4945269.1 | *F. gaditjirri* | *Nisikadoi* | Not known | 91% | 76.19 |
| KAG5655603.1 | *F. avenaceum* | *Tricinctum* | Plant pathogen | 91% | 76.19 |
| XP_018760148.1 | *F. verticillioides* 7600 | *Fujikuroi* | Plant pathogen | 91% | 75.76 |
| XP_031031185.1 | *F. oxysporum* Fo47 | *Oxysporum* | Plant endophytic | 91% | 75.76 |
| RYC87346.1 | *F. oxysporum* f. sp. *narcissi* | *Oxysporum* | Plant pathogen | 91% | 75.76 |
| EGU81637.1 | *F. oxysporum* f. sp. *conglutinans* Fo5176 | *Oxysporum* | Plant pathogen | 91% | 75.76 |
| KAH7245353.1 | *F. tricinctum* | *Tricinctum* | Plant pathogen | 99% | 75.6 |
| KAF4998722.1 | *F. graminum* | *Heterosporum* | Not known | 100% | 75.4 |
| SCV32335.1 | *F. fujikuroi* | *Fujikuroi* | Plant pathogen | 91% | 75.32 |
| RBR01797.1 | *F. verticillioides* | *Fujikuroi* | Plant pathogen | 91% | 75.32 |
| XP_018252424.1 | *F. oxysporum* f. sp. *lycopersici* 4287 | *Oxysporum* | Plant pathogen | 91% | 75.32 |
| KAF4341936.1 | *F. beomiforme* | *Babinda* | not known | 91% | 74.89 |
| KAG5750526.1 | *F. xylarioides* | *Fujikuroi* | Plant pathogen | 91% | 74.89 |
| XP_041677642.1 | *F. mangiferae* | *Fujikuroi* | Plant pathogen | 91% | 74.89 |
| KAF5689528.1 | *F. circinatum* | *Fujikuroi* | Plant pathogen | 91% | 74.89 |
| XP_036533940.1 | *F. subglutinans* | *Fujikuroi* | Plant pathogen | 91% | 74.46 |
| SPJ76027.1 | *F. torulosum* | *Tricinctum* | Not known | 99% | 74.4 |
| KAG5659696.1 | *F. avenaceum* | *Tricinctum* | Plant pathogen | 99% | 74.4 |
| KAF5667294.1 | *F. heterosporum* | *Heterosporum* | hyperparasite of *Claviceps purpurea* | 100% | 73.81 |
| KAF4947933.1 | *F. sarcochroum* | *Lateritium* | Not known | 99% | 71.37 |
| KAJ4252383.1 | *F. torreyae* | No complex assigned | Plant pathogen | 95% | 70.04 |

<sup>1</sup> The presence of a secretion signal and a chloroplast transit peptide were predicted using SignalP v6.0 (Teufel et al. 2022) and LOCALIZER v1.0 (Sperschneider et al. 2017), respectively. All proteins showed probability values above 0.99 and 0.95 for the presence of a secretion signal and chloroplast transit peptide sequences, respectively.
<sup>2</sup> *Fusarium* species were assigned to the different *Fusarium* species complexes based on the classification done by Waalwijk et al (2018).

**Supplementary Table S3.**
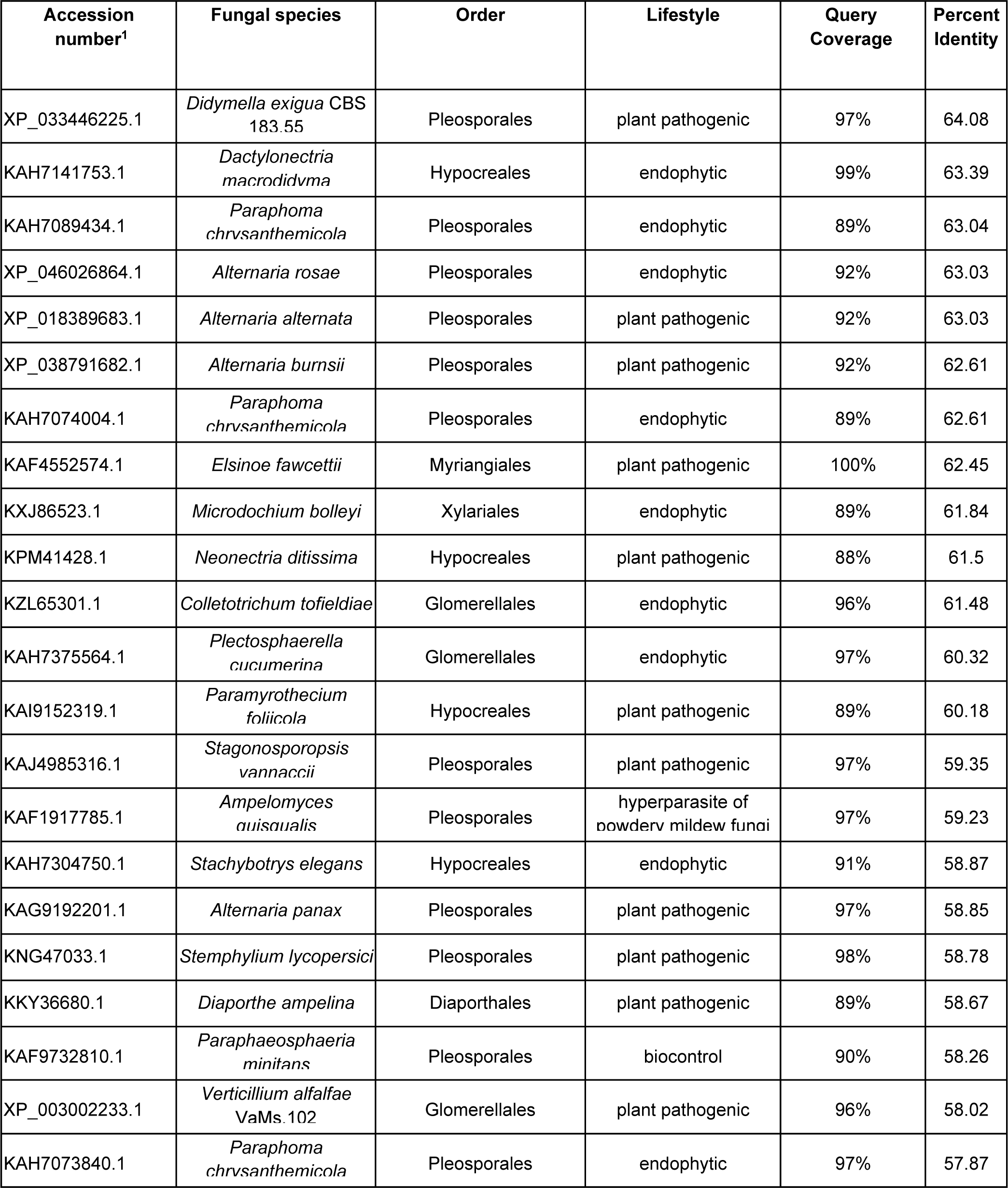
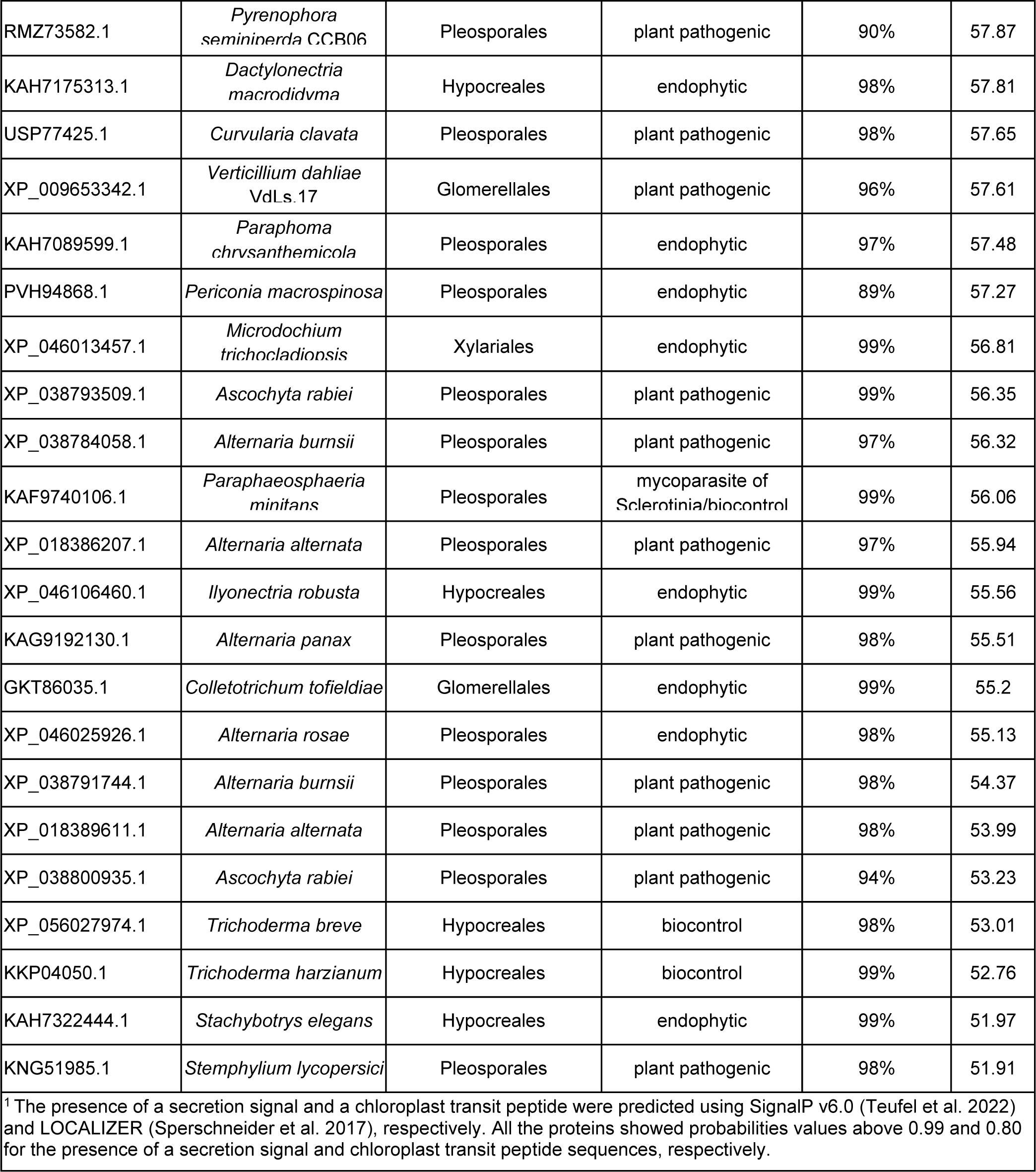
FgTPP1 protein orthologs in different genera of the *Ascomycota* phylum.

**Supplementary Table S4.**
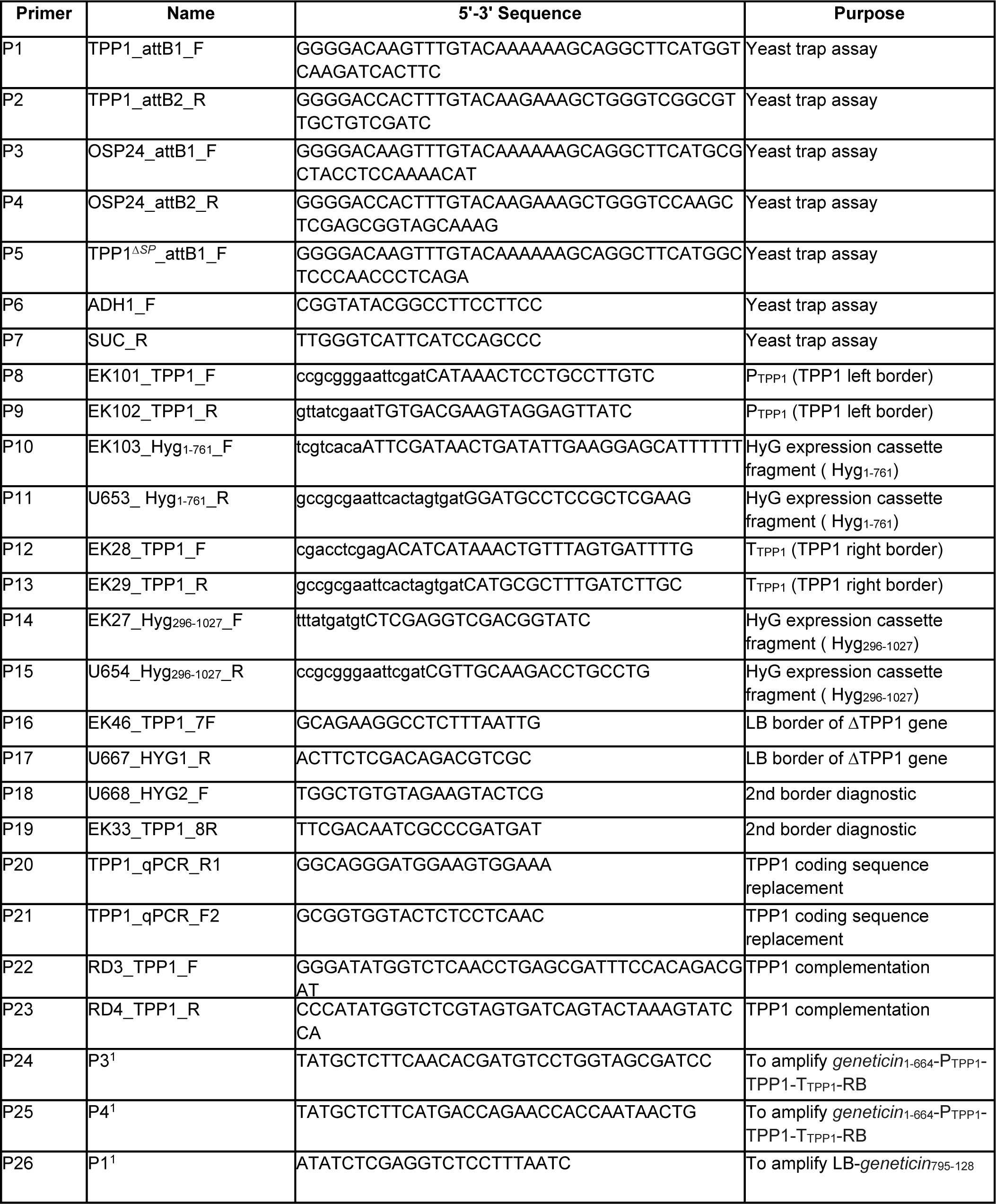
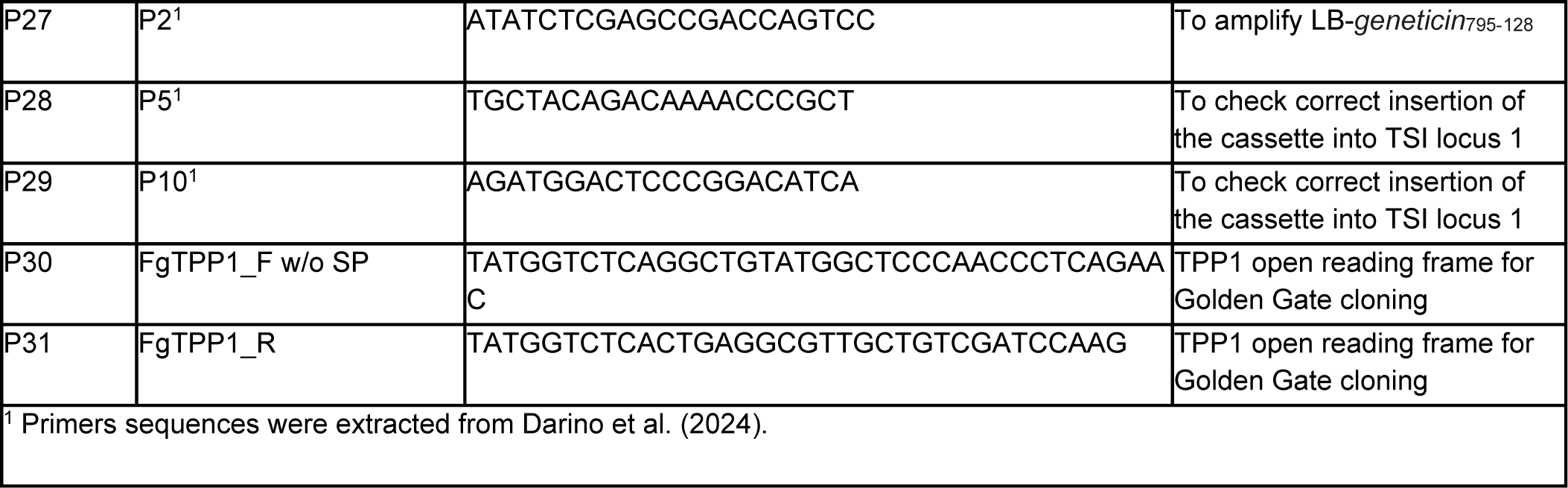
List of primers used in this work.

